# On the generalizability of diffusion MRI signal representations across acquisition parameters, sequences and tissue types: chronicles of the MEMENTO challenge

**DOI:** 10.1101/2021.03.02.433228

**Authors:** Alberto De Luca, Andrada Ianus, Alexander Leemans, Marco Palombo, Noam Shemesh, Hui Zhang, Daniel C Alexander, Markus Nilsson, Martijn Froeling, Geert-Jan Biessels, Mauro Zucchelli, Matteo Frigo, Enes Albay, Sara Sedlar, Abib Alimi, Samuel Deslauriers-Gauthier, Rachid Deriche, Rutger Fick, Maryam Afzali, Tomasz Pieciak, Fabian Bogusz, Santiago Aja-Fernández, Evren Özarslan, Derek K Jones, Haoze Chen, Mingwu Jin, Zhijie Zhang, Fengxiang Wang, Vishwesh Nath, Prasanna Parvathaneni, Jan Morez, Jan Sijbers, Ben Jeurissen, Shreyas Fadnavis, Stefan Endres, Ariel Rokem, Eleftherios Garyfallidis, Irina Sanchez, Vesna Prchkovska, Paulo Rodrigues, Bennet A Landman, Kurt G Schilling

**Affiliations:** PROVIDI Lab, Image Sciences Institute, University Medical Center Utrecht, Utrecht, the Netherlands; Department of Neurology, UMC Utrecht Brain Center, University Medical Center Utrecht, Utrecht, the Netherlands; Champalimaud Research, Champalimaud Centre for the Unknown, Lisbon, Portugal; Centre for Medical Image Computing, Department of Computer Science, University College London, London, United Kingdom; Clinical Sciences Lund, Radiology, Lund University, Lund, Sweden; Department of Radiology, University Medical Center Utrecht; Inria Sophia Antipolis – Méditerranée, Université Côte d’Azur, Sophia Antipolis, France; Istanbul Technical University, Istanbul, Turkey; TRIBVN Healthcare, Paris, France; Cardiff University Brain Research, Imaging Centre (CUBRIC), School of Psychology, Cardiff University, Cardiff, United Kingdom; AGH University of Science and Technology, Kraków, Poland; LPI, ETSI Telecomunicación, Universidad de Valladolid, Valladolid, Spain; Department of Biomedical Engineering, Linköping University, Linköping, Sweden; Center for Medical Image Science and Visualization, Linköping University, Linköping, Sweden; School of Instruments and Electronics, North University of China, Taiyuan, China; Department of Physics, University of Texas at Arlington, Arlington, USA; NVIDIA Corporation, Bethesda, USA; National Institute of Health, Bethesda, USA; Imec-Vision lab, Department of Physics, University of Antwerp, Antwerp, Belgium; Intelligent Systems Engineering, Indiana University Bloomington, Indiana, USA; Leibniz Institute for Materials Engineering — IWT, Faculty of Production Engineering, University of Bremen, Bremen, Germany; Department of Psychology and the eScience Institute, University of Washington, Seattle, WA USA; QMENTA Inc, Boston, USA; Vanderbilt University Institute of Imaging Science, Vanderbilt University, Nashville, USA; Department of Radiology and Radiological Science, Vanderbilt University Medical Center, Nashville, USA

## Abstract

Diffusion MRI (dMRI) has become an invaluable tool to assess the microstructural organization of brain tissue. Depending on the specific acquisition settings, the dMRI signal encodes specific properties of the underlying diffusion process. In the last two decades, several signal representations have been proposed to fit the dMRI signal and decode such properties. Most methods, however, are tested and developed on a limited amount of data, and their applicability to other acquisition schemes remains unknown. With this work, we aimed to shed light on the generalizability of existing dMRI signal representations to different diffusion encoding parameters and brain tissue types. To this end, we organized a community challenge - named MEMENTO, making available the same datasets for fair comparisons across algorithms and techniques. We considered two state-of-the-art diffusion datasets, including single-diffusion-encoding (SDE) spin-echo data from a human brain with over 3820 unique diffusion weightings (the MASSIVE dataset), and double (oscillating) diffusion encoding data (DDE/DODE) of a mouse brain including over 2520 unique data points. A subset of the data sampled in 5 different voxels was openly distributed, and the challenge participants were asked to predict the remaining part of the data. After one year, eight participant teams submitted a total of 80 signal fits. For each submission, we evaluated the mean squared error, the variance of the prediction error and the Bayesian information criteria. Most predictions predicted either multi-shell SDE data (37%) or DODE data (22%), followed by cartesian SDE data (19%) and DDE (18%). Most submissions predicted the signals measured with SDE remarkably well, with the exception of low and very strong diffusion weightings. The prediction of DDE and DODE data seemed more challenging, likely because none of the submissions explicitly accounted for diffusion time and frequency. Next to the choice of the model, decisions on fit procedure and hyperparameters play a major role in the prediction performance, highlighting the importance of optimizing and reporting such choices. This work is a community effort to highlight strength and limitations of the field at representing dMRI acquired with trending encoding schemes, gaining insights into how different models generalize to different tissue types and fiber configurations over a large range of diffusion encodings.

## Introduction

Diffusion Magnetic Resonance Imaging (dMRI) is a powerful tool to investigate microstructural properties of biologic tissues in-vivo (A. L. Alexander et al. 2007; J. D. Tournier, Mori, and Leemans 2011) with applications in neuroimaging studying brain development (Ouyang et al. 2019), plasticity (Blumenfeld-Katzir et al. 2011), aging (Baker et al. 2014), as well as changes upon disease for diagnostic and monitoring purposes in various conditions such as Alzheimer’s disease (Doan et al. 2017; Weston et al. 2015), multiple sclerosis (Inglese and Bester 2010; De Santis et al. 2019), Parkinson’s disease (Atkinson-Clement et al. 2017), brain tumours (Costabile et al. 2019), etc. The signal measured in dMRI is sensitized to the microscopic motion of water molecules, which is hindered and restricted by the presence of biologic membranes, thus carrying information about the cellular organization. Over the last decade, an increasing number of techniques have been proposed in the literature to describe the dMRI signal and provide biomarkers of tissue microstructure and have been recently complemented with various machine learning approaches.(D. C. Alexander et al. 2017; Ghosh, Ianus, and Alexander 2018; D. S. Novikov et al. 2019; Poulin et al. 2019; Ravi et al. 2019)

The standard acquisition strategy for dMRI data is single diffusion encoding (SDE), which employs a pair of diffusion weighting gradients with identical areas, usually embedded before and after the refocusing pulse in a spin echo preparation, a sequence widely known also as pulsed gradient spin-echo (Stejskal and Tanner 1965). The SDE sequences are characterized by the gradient strength (G), duration (δ), time interval between the onset of the two gradients (Δ) and gradient orientation (*ĝ*). The scalar parameters (G, δ, Δ) are usually combined to describe the diffusion weighting of the sequence, also referred to as the b-value. For SDE sequences, b = γ^2^G^2^δ^2^(Δ- δ/3), where γ is the gyromagnetic ratio. In the majority of SDE acquisitions δ and Δ are fixed and G is varied to change the b-value, although varying the gradient duration and diffusion time can provide additional orthogonal measurements (D. S. Novikov et al. 2019). Over the last decade, diffusion sequences which further vary the gradient waveform within one measurement, such as double diffusion encoding (DDE) (Mitra 1995; Henriques et al. 2020) or b-tensor encoding approaches (Lasic et al. 2014; C. F. Westin et al. 2016), have been gaining interest as they can further improve the specificity of the measurements towards the underlying tissue microstructure. Other approaches replace the pulsed gradients with oscillating gradient waveforms to probe diffusion on a range of (shorter) time scales (Does, Parsons, and Gore 2003; Burcaw, Fieremans, and Novikov 2015), measurements which can also be performed with different gradient orientations in a double oscillating diffusion encoding (DODE) fashion (Ianus et al. 2017, 2018). While the majority of recent dMRI studies employ SDE sequences, such advanced acquisitions are steadily gaining popularity.

The most widely used dMRI technique for brain imaging in the clinic is diffusion tensor imaging (DTI)(Basser, Mattiello, and LeBihan 1994), which assumes that water diffusion in the underlying tissue can be described by a Gaussian anisotropic process. As minimum requirements, the tensor parameters can be estimated from SDE sequences with a single b-value (usually about 1000 s/mm^2^) and at least 6 non-collinear directions in addition to non-diffusion weighted data (b = 0 s/mm^2^). Although simple and robust, DTI cannot describe the signal decay in correspondence of higher b-values (e.g., above about 1500 s/mm^2^ in the living brain) and cannot distinguish between multiple fibre populations, for instance in areas of crossing fibres (B. Jeurissen et al. 2013). After DTI, a plethora of techniques have been introduced to capture the dMRI signal decay over a wider range of parameter values (D. C. Alexander et al. 2017; D. S. Novikov et al. 2019).

dMRI models can generally be regarded as biophysical models, signal representations or somewhere in between (Ghosh, Ianus, and Alexander 2018; Ileana O. Jelescu and Budde 2017). Biophysical models usually employ multiple water compartments to describe the dMRI signal in the tissue in order to capture microscopic metrics such as intracellular signal fraction, cell size, shape etc (Stanisz et al. 1997; Jespersen et al. 2007; Y. Assaf et al. 2008; Daniel C. Alexander 2008; E. Fieremans et al. 2013; Palombo et al. 2020; Panagiotaki et al. 2012; Fan et al. 2020; Zhang et al. 2012). Several biophysical models have been proposed in the literature and vary in terms of the number of compartments, diffusion model (hindered/restricted), number of fibre populations, fibre orientation distributions, etc. Signal representations on the other hand, usually provide a statistical description to capture the signal decay without explicitly modelling the underlying tissue composition (Yablonskiy, Bretthorst, and Ackerman 2003; J. H. Jensen et al. 2005; Ozarslan et al. 2009; Els Fieremans, Jensen, and Helpern 2011; Steven, Zhuo, and Melhem 2014; Ozarslan et al. 2013). Next to these two main families, there are also hybrid approaches that, for instance, aim to characterize the fibre orientation distribution without explicitly modelling the fibre composition (J. D. Tournier et al. 2008; Ben Jeurissen et al. 2014), or use a statistical model for different compartments (Scherrer et al. 2016; Pasternak et al. 2009; De Luca, Bertoldo, and Froeling 2017), which can be defined a-priori or driven from the data (Keil et al. 2017; De Luca et al. 2018). Besides these “classical” approaches to model dMRI, the last couple of years have witnessed a vast increase in the number of machine learning techniques applied to dMRI to predict signal decay (Golkov et al. 2016; Grussu et al. 2020), fibre orientations (Poulin et al. 2019; Nath, Schilling, et al. 2019) or the underlying tissue parameters (Nedjati-Gilani et al. 2017).

The choice of dMRI technique depends on many factors, such as the purpose of the experiment, the amount and quality of the data, the number and strength of b-values, angular resolution, etc, and generally no consensus has been reached on what a state-of-the-art diffusion experiment should include. Nevertheless, for a given acquisition, comparing diffusion models can provide valuable information about which approaches best describe the signal and can be generalized to predict measurements outside the initial range. In the literature, there have been various studies which aimed to compare brain tissue models in terms of goodness of fit and signal prediction, with an emphasis on white matter (WM) (Panagiotaki et al. 2012; U. Ferizi et al. 2015; I. O. Jelescu et al. 2014; Rokem et al. 2015) and less on gray matter (GM)(Yaniv Assaf 2019). However, such studies usually focused on a certain group of models, for instance multi-compartment biophysical models were investigated in (Panagiotaki et al. 2012; U. Ferizi et al. 2015), while (Wang et al. 2017) looked in more detail at signal representations.

Open challenges play an important role to gain a better understanding of how various models capture the dMRI signal decay, as they put forward rich datasets and well-defined tasks and usually receive submissions from across the modelling landscapes (Uran Ferizi et al. 2017; Schilling et al. 2019; Pizzolato et al. 2020). The last diffusion microstructure challenge which included a comprehensive dMRI acquisition (Uran Ferizi et al. 2017) was organized in 2015 and focused on modelling the dMRI signal acquired on the Connectome scanner for two ROIs in white matter: genu of the Corpus Callosum with mostly aligned fibres and fornix with a more complex fibre configuration. The challenge included a rich dataset acquired with many combinations of gradient strengths, durations and diffusion times and the goal was to predict unseen shells with parameter values within the range used for the provided data. Since the end of this challenge, many novel approaches have been proposed, including a booming application of machine learning techniques for data fitting and prediction (Golkov et al. 2016; Nedjati-Gilani et al. 2017; Nath, Schilling, et al. 2019; Ravi et al. 2019; Poulin et al. 2019). Moreover, previous challenges (Uran Ferizi et al. 2017; Schilling et al. 2019; Pizzolato et al. 2020) included only diffusion data acquired with standard SDE sequences, and do not provide any insight into the different approaches available to analyse advanced sequences such as DDE.

In this challenge we set to evaluate the ability of different dMRI modeling approaches to capture the dMRI signal contrast from state-of-the-art acquisitions performed with SDE, including shells with an order of magnitude higher angular resolution compared to previous challenges (Uran Ferizi et al. 2017; Schilling et al. 2019; Pizzolato et al. 2020), as well as DDE and DODE data. Further, we aim to investigate the relationship between the goodness of fit and tissue type, acquisition parameters, and diffusion sensitization. Finally, this challenge acts as a benchmark database for the evaluation of future models focusing on healthy tissue as the full datasets are made available (https://github.com/PROVIDI-Lab/MEMENTO.git).

## Methods

Section 2.1 presents an overview of the data that was used in the MEMENTO challenge and of the reasoning behind the selection of specific brain locations. A description of the methods used in the received submissions is reported in section 2.2, whereas section 2.3 illustrates the analyses we performed on the collected signal predictions.

### Challenge data

#### MRI acquisition

The main aim of the MEMENTO challenge was to investigate how well existing models can represent the dMRI signal collected with i) different gradient encoding schemes, and ii) from different tissue types.

To investigate how the existing models can predict data sampled with different diffusion sensitization, we selected two datasets containing extensively sampled dMRI brain data with 4 encoding schemes: multi-shell SDE (SDE-MS), SDE with gradients sampled in a cartesian grid (SDE-GRID), DDE and DODE. The SDE acquisitions were performed in a healthy volunteer with a 3T scanner as part of the MASSIVE datasets (Froeling et al. 2017a), a collection of 18 MRI sessions containing unique dMRI data performed with a 3T scanner (Philips Healthcare, The Netherlands) with voxel-size 2.5mm^3^ isotropic, echo time TE=100ms and repetition time TR between 7 and 7.5s. The DDE and DODE data were sampled from an ex-vivo mouse brain imaged with a 16.4T scanner (Bruker) with imaging resolution 0.12×0.12×0.7mm^3^, TE=52ms, TR=3s (Ianus et al. 2018). The diffusion parameters of the acquired data are reported in Table 1 and Table 2. To establish the signal to noise ratio (SNR) of the data, we considered the datapoints collected at b = 0 s/mm2, removed eventual outliers and defined the average SNR as the ratio between the average non-weighted value and its standard deviation. A point was defined as an outlier when its value was outside the confidence interval defined by the median value ± 2 times the robust standard deviation of the data (see (Chang, Jones, and Pierpaoli 2005)). The SNR of the 5 selected signals at b = 0 s/mm2 was 15 ± 3 for SDE-MS, 16 ± 3 for SDE-GRID, 76 ± 37 for DDE and 66 ± 28 for DODE.

**Table 1 -.** A description of the SDE encoding acquired as part of the MASSIVE data, and their subdivision in training and evaluation data. No data points of SDE-MS at b = 4000 s/mm^2^ and SDE-GRID at b > 7600 s/mm^2^ were provided for training. The ratio between training and evaluation data was about 1:3 for SDE-MS and 3:5 for SDE-GRID.

| Table 1 - A description of the SDE encoding acquired as part of the MASSIVE data, and their subdivision in training and evaluation data. No data points of SDE-MS at $b = 4000 \text{ s/mm}^2$ and SDE-GRID at $b > 7600 \text{ s/mm}^2$ were provided for training. The ratio between training and evaluation data was about 1:3 for SDE-MS and 3:5 for SDE-GRID. | | | |
| --- | --- | --- | --- |
| Diffusion Encoding | Number of directions | Training | Evaluation |
| **SDE-MS data of a healthy volunteer acquired at 3T in 18 sessions (13 shells)** |  |  |  |
| $b = 0 \text{ s/mm}^2$ | 430 | 20 | 410 |
| $b = 5 \text{ s/mm}^2$ | 30 | 8 | 22 |
| $b = 10 \text{ s/mm}^2$ | 30 | 7 | 23 |
| $b = 25 \text{ s/mm}^2$ | 40 | 10 | 30 |
| $b = 40 \text{ s/mm}^2$ | 40 | 10 | 30 |
| $b = 60 \text{ s/mm}^2$ | 20 | 5 | 15 |
| $b = 80 \text{ s/mm}^2$ | 20 | 5 | 15 |
| $b = 140 \text{ s/mm}^2$ | 20 | 5 | 15 |
| $b = 250 \text{ s/mm}^2$ | 30 | 8 | 22 |
| $b = 500 \text{ s/mm}^2$ | 250 | 62 | 188 |
| $b = 1000 \text{ s/mm}^2$ | 500 | 125 | 375 |
| $b = 2000 \text{ s/mm}^2$ | 500 | 125 | 375 |
| $b = 3000 \text{ s/mm}^2$ | 500 | 125 | 375 |
| $b = 4000 \text{ s/mm}^2$ | 600 | 0 | 600 |
| Total | 3010 | 515 | 2495 |
| **SDE-GRID data of a healthy volunteer acquired at 3T in 18 sessions** |  |  |  |
| $b = 0 \text{ s/mm}^2$ | 430 | 20 | 410 |
| b = 141-563 s/mm<sup>2</sup> | 58 | 22 | 36 |
| b = 750-938 s/mm<sup>2</sup> | 54 | 22 | 32 |
| b = 1125-1875 s/mm<sup>2</sup> | 170 | 72 | 98 |
| b = 2016-2766 s/mm<sup>2</sup> | 248 | 102 | 146 |
| b = 3000-3938 s/mm<sup>2</sup> | 314 | 129 | 185 |
| b = 4125-4875 s/mm<sup>2</sup> | 168 | 68 | 100 |
| b = 5016-6891 s/mm<sup>2</sup> | 196 | 65 | 131 |
| b = 7687-9000 s/mm<sup>2</sup> | 32 | 0 | 32 |
| Total | 1670 | 500 | 1170 |

**Table 2 -.** The table describes the subdivision in training and evaluation data of the unique combinations of diffusion weighting b and diffusion time (Δ) / oscillation frequency (*f*) for the DDE and DODE data, respectively. In the DDE training set, the data acquired with Δ = 5ms was provided with the exception of the largest diffusion weighting (b = 4000 s/mm^2^), and no data acquired with Δ = 10ms was provided for training. The mixing time for DDE sequences was 18.3ms. For DODE, the data points acquired with the three lower oscillation frequencies (66, 100, 133 Hz) were provided with the exception of b = 4000 s/mm^2^, and no data acquired with f = 166 and 200 Hz was provided for training. The separation time between gradient waveforms was 5ms. For DDE and DODE data, the sequence description included information about gradient strengths, directions, timing parameters as well as the elements of the B-matrix.

| Diffusion Encoding | Number of directions | Training | Evaluation |
| --- | --- | --- | --- |
| **DDE data from a mouse brain ex-vivo acquired at 16.4T** |  |  |  |
| $\Delta=5\text{ms}$, $b = 0\text{ s/mm}^2$ | **40** | **32** | **8** |
| $\Delta=5\text{ms}$, $b = 1000\text{ s/mm}^2$ | **72** | **72** | **0** |
| $\Delta=5\text{ms}$, $b = 1750\text{ s/mm}^2$ | **72** | **72** | **0** |
| $\Delta=5\text{ms}$, $b = 2500\text{ s/mm}^2$ | **72** | **72** | **0** |
| $\Delta=5\text{ms}$, $b = 3250\text{ s/mm}^2$ | **72** | **72** | **0** |
| $\Delta=5\text{ms}$, $b = 4000\text{ s/mm}^2$ | **72** | **0** | **72** |
| $\Delta=10\text{ms}$, $b = 0\text{ s/mm}^2$ | **40** | **0** | **40** |
| $\Delta=10\text{ms}$, $b = 1000\text{ s/mm}^2$ | **72** | **0** | **72** |
| $\Delta=10\text{ms}$, $b = 1750\text{ s/mm}^2$ | **72** | **0** | **72** |
| $\Delta=10\text{ms}$, $b = 2500 \text{ s/mm}^2$ | **72** | **0** | **72** |
| $\Delta=10\text{ms}$, $b = 3250 \text{ s/mm}^2$ | **72** | **0** | **72** |
| $\Delta=10\text{ms}$, $b = 4000 \text{ s/mm}^2$ | **72** | **0** | **72** |
| Total | **800** | **320** | **480** |
| **DODE data from a mouse brain ex-vivo acquired at 16.4T** |  |  |  |
| $f = 66, 100, 133\text{Hz}$<br>$b = 0 \text{ s/mm}^2$ | **3x40** | **3x32** | **3x8** |
| $f = 66, 100, 133\text{Hz}$<br>$b = 1000 \text{ s/mm}^2$ | **3x72** | **3x72** | **0** |
| $f = 66, 100, 133\text{Hz}$<br>$b = 1750 \text{ s/mm}^2$ | **3x72** | **3x72** | **0** |
| $f = 66, 100, 133\text{Hz}$<br>$b = 2500 \text{ s/mm}^2$ | **3x72** | **3x72** | **0** |
| $f = 66, 100, 133\text{Hz}$<br>$b = 3250 \text{ s/mm}^2$ | **3x72** | **3x72** | **0** |
| $f = 66, 100, 133\text{Hz}$<br>$b = 4000 \text{ s/mm}^2$ | **3x72** | **0** | **3x72** |
| $f = 166, 200\text{Hz}$<br>$b = 0 \text{ s/mm}^2$ | **2x40** | **0** | **2x40** |
| $f = 166, 200\text{Hz}$<br>$b = 1000 \text{ s/mm}^2$ | **2x72** | **0** | **2x72** |
| $f = 166, 200\text{Hz}$<br>$b = 1750 \text{ s/mm}^2$ | **2x72** | **0** | **2x72** |
| $f = 166, 200\text{Hz}$<br>$b = 2500 \text{ s/mm}^2$ | **2x72** | **0** | **2x72** |
| $f = 166, 200\text{Hz}$<br>$b = 3250 \text{ s/mm}^2$ | **2x72** | **0** | **2x72** |
| $f = 166, 200\text{Hz}$<br>$b = 4000 \text{ s/mm}^2$ | **2x72** | **0** | **2x72** |
| Total | **2000** | **960** | **1040** |

#### Signals selection

Five signals were selected for each dataset from brain voxels exhibiting different microstructural organization. For the human MASSIVE dataset, the voxels aimed to include WM signals with an increasing number of crossing fiber configurations from 1 to 3, deep gray matter (DGM) and cortical gray matter (CGM) and the selection was based on visual inspection of the fiber orientation distribution (FOD). The FOD was derived with constrained spherical deconvolution (J.-D. Tournier, Calamante, and Connelly 2007) of data at b = 0, 3000 s/mm^2^ (500 directions) using a recursively calibrated response function(Tax et al. 2014) approach implemented in ExploreDTI. Given the relatively large imaging resolution of the MASSIVE dataset, 2.5mm^3^ isotropic, care was taken in selecting voxels not located at tissue interfaces, also using the FOD as guidance. For the mouse brain, the five voxels were placed in white matter tracts with different microstructures and based on visual comparison with an anatomical atlas, as the number of collected gradient orientations per diffusion weighting was insufficient to reliably estimate the FOD. Specifically, the five voxels were placed in medial corpus callosum, lateral corpus callosum, internal capsule, a fanning region of the internal capsule and the fimbria, respectively. An illustration of the locations and tissue types of the selected voxels is shown in Figure 1.

**Figure 1:**
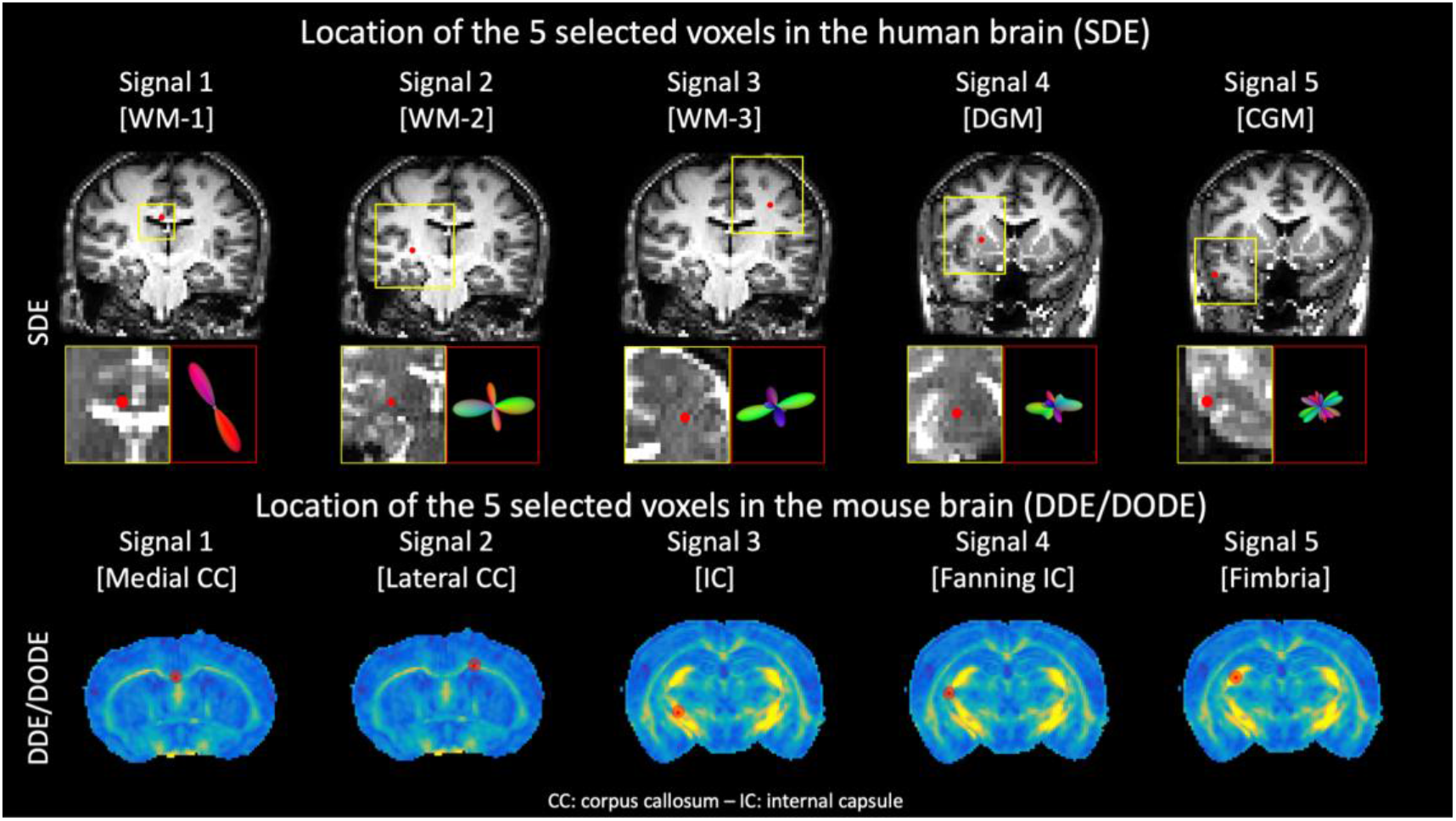
The locations of the 5 voxels selected for the SDE (top) and DDE/DODE data (bottom). The SDE data are sampled from a human brain scanned at 3T with imaging resolution 2.5mm^3^ isotropic, and is part of the MASSIVE dataset. The 5 locations were chosen in 5 different tissue types, such as white matter (WM) with increasing fiber complexity (signals 1-3, as exemplified by the shown fiber orientation distribution), deep gray matter (DGM) and cortical gray matter (CGM). DDE and DODE were sampled in a mouse brain at 16.4T with imaging resolution 0.12×0.12×0.7mm^3^. The selected signals are all taken from WM locations with well-established differences in fiber organization, as shown by the red dots overlaid on the fractional anisotropy map.

#### Training and evaluation data

The 20 selected measurement sets (5 signal locations x 4 diffusion encodings) were subdivided in training and evaluation data. The challenge participants were provided with the training data and the corresponding diffusion encoding information, and asked to predict the evaluation data. The proportion of training and evaluation data was not constant among data encodings. For SDE, the training data consisted of about 500 gradient directions uniformly subsampled from all available data except for the largest diffusion weightings, which corresponds to about 17% of SDE-MS, and 30% of SDE-GRID. For DDE and DODE, we provided a larger amount of training data to take into account the need to model an additional encoding dimension (e.g., time and frequency), respectively 40% and 48%. To evaluate the ability of the tested models to predict unseen data points, all the data corresponding to specific diffusion weightings was removed from the training data, as reported in Table 1 and Table 2.

### Signal predictions

We received initial submissions from 9 teams, but 2 of the 9 teams did not provide valid submissions and were not included in this analysis. The remaining 7 teams submitted a total of 80 valid signal predictions that were considered in the subsequent analyses. Of these, 31 submissions predicted the SDE-MS signals (37%), 16 the SDE-GRID signals (19%), 15 the DDE signals (18%) and 18 the DODE signals (22%). When a model was applied more than once to predict a given set of signals, only the submissions corresponding to the best and worst prediction were analyzed, to simplify the presentation and interpretation of the results. The final selection of predictions included in this analysis is reported in Table 3. When multiple predictions with a given model were submitted, the best and worst predictions were identified by adding the labels “_best” and “_worst” to the model name.

**Table 3:** The valid signal predictions submitted to the MEMENTO challenge. For each method, we report the acronym and the main reference, the “category”, special notes on the fit procedure, and the data it has been applied to. The following predictions were subdivided in the following categories: tensor-based (TENS), multi-compartment model (MCM), parametric representation (PAR), deep learning-based (DL).

| Model name | Category | Notes | SDE-MS | SDE-GRID | DDE | DODE |
| --- | --- | --- | --- | --- | --- | --- |
| DTI (Basser, Mattiello, and LeBihan 1994) | TENS | Linear-Least Squares | X | X | X | X |
| DKI (J. H. Jensen et al. 2005) | TENS | Weighted-Least Squares | X | X | X | X |
| DKI+Offset<br>(Morez et al.<br>2020) | TENS | Constrained<br>Non-Linear fit | X | X | X |  |
| DTD-cov (C. F.<br>Westin et al.<br>2016) | TENS | Constrained<br>Non-Linear fit |  |  | X | X |
| DTD-cov (C. F.<br>Westin et al.<br>2016) + Offset | TENS | Constrained<br>Non-Linear fit |  |  | X | X |
| Ball&Stick<br>(Behrens et al.<br>2003) | MCM | Implemented in<br>Dmipy (Fick,<br>Wassermann,<br>and Deriche<br>2019) | X |  |  |  |
| Ball&Racket<br>(Sotiropoulos,<br>Behrens, and<br>Jbabdi 2012) | MCM | Implemented in<br>Dmipy (Fick,<br>Wassermann,<br>and Deriche<br>2019) | X |  |  |  |
| NODDI-Watson<br>(Zhang et al.<br>2012) | MCM | Implemented in<br>Dmipy (Fick,<br>Wassermann,<br>and Deriche<br>2019) | X |  |  |  |
| NODDI -<br>Bingham (Tariq<br>et al. 2016) | MCM | Implemented in<br>Dmipy (Fick,<br>Wassermann,<br>and Deriche<br>2019) | X |  |  |  |
| SMT (Kaden et<br>al. 2016) | MCM | Implemented in<br>Dmipy (Fick,<br>Wassermann,<br>and Deriche<br>2019) | X |  |  |  |
| NODDI-SMT | MCM | Implemented in<br>Dmipy (Fick,<br>Wassermann,<br>and Deriche<br>2019) | X |  |  |  |
| MCMDI (Kaden<br>et al. 2016) | MCM | Implemented in<br>Dmipy (Fick,<br>Wassermann,<br>and Deriche<br>2019) | X |  |  |  |
| ActiveAx (D. C.<br>Alexander et al.<br>2010) | MCM | Implemented in<br>Dmipy (Fick,<br>Wassermann,<br>and Deriche<br>2019) | X |  |  |  |
| SHORE<br>(Ozarslan et al.<br>2009) | PAR | From<br>DeepSHORE<br>(Nath, Lyu, et<br>al. 2019) | X | X | X | X |
| MAP-MRI<br>(Fick et al.<br>2016) | PAR | Implemented in<br>Dmipy (Fick,<br>Wassermann, and Deriche 2019) | X |  |  |  |
| MAP-MRI+Reg (Fick et al. 2016) | PAR | Implemented in Dipy (Garyfallidis et al. 2014) | X | X |  |  |
| NeuralNet | DL | Perceptron 1 Layer 50 nodes | X | X | X | X |
| NeuralNet+Rein f (Williams 1992) | DL | Perceptron 7 Layers optimized with NAS (Zoph and Le 2016) | X | X | X | X |

To follow, we present an overview of the submissions we received grouped in four different categories.

#### Tensor and beyond

Diffusion tensor imaging (DTI, Basser et al. 1994) is one of the most common quantification methods for dMRI data acquired with at least one diffusion-weighting and 6+ gradient directions. DTI is based on the three-dimensional generalization of the seminal works of Stejskal and Tanner (Stejskal and Tanner 1965), and assumes the diffusion process to be Gaussian (i.e., not restricted). In the living brain, such assumption is typically satisfied when collecting dMRI data with diffusion weightings in the range b = 800-1200 s/mm^2^. While DTI typically does not accurately characterize complex diffusion environments where, for example, multiple diffusion mode (e.g., tissue diffusion vs blood pseudo-diffusion (Le Bihan et al. 1988)) or “crossing-fibers” co-exist (Wedeen et al. 2005), it is one of the most common dMRI signal representations in clinical application, especially thanks to its sensitivity to microstructural changes in health and pathology. Keeping in mind all of the above, the DTI method was applied in this work to SDE-MS, DDE and DODE data to serve as baseline reference using a weighted-least-squares fit.

In 2005, Jensen and colleagues introduced the diffusion kurtosis imaging (DKI) method (J. H. Jensen et al. 2005), an extension of DTI that allows to account for and quantify the amount of non-Gaussian diffusion that is observed at stronger diffusion weightings (e.g., b > 1400 s/mm^2^ in the living brain). The DKI model requires the collection of at least 21 unique measurements, including 2 non-zero diffusion-weightings and 15+ unique gradient directions, and allows to quantify the amount of excess kurtosis of the diffusion process. While DKI is suitable for dMRI data acquired with a stronger diffusion weighting than DTI, nevertheless, there is a theoretical maximum to the diffusion weighting that can be fit (J. H. Jensen et al. 2005; Jens H. Jensen and Helpern 2010), and most brain applications use a maximum b-value equal to b = 3000 s/mm^2^. Despite these theoretical limits, DKI was fit to all data included in this work using a constrained non-linear least-squares fit enforcing positivity in diffusion and kurtosis metrics and a monotonic decay of the signal. When fitting data acquired with strong diffusion weighting, it might be beneficial to take into account the presence of a minimum signal offset due to Rician noise in the measurements (Gudbjartsson and Patz 1995; Basu, Fletcher, and Whitaker 2006). One of the submissions considered in this work extended the DKI method with an offset term to account for such effect (DKI+Offset)(Morez et al. 2020). This method was fit to SDE data by extending the classic DKI model with an additional degrees of freedom (22 + 1 = 23 free parameters), and to DDE and DODE data by extending a fourth order covariance tensor (28 + 1 = 29 free parameters) (C.-F. Westin et al. 2016).

#### Multicompartment models

Multi-compartment models are a family of methods that allow to model the dMRI signal by means of biophysical features (Panagiotaki et al. 2012; Ileana O. Jelescu and Budde 2017). The assumption behind these models is that the dMRI signal acquired in a voxel can be described as the linear combination of the signal profiles of each component that is present in a specific voxel.

All the submissions we received based on multicompartment models were computed with custom implementations of previously introduced methods with the “Diffusion Microstructure Imaging in Python” toolbox (dmipy). The submissions considered only SDE data, and were based on three basic components: the intra-axonal compartment was modelled as a stick or a cylinder, whereas the extra-axonal anisotropic compartment was modelled as a zeppelin (axially symmetric tensor) and the cerebrospinal fluid contribution modelled as isotropic diffusion (sphere). Depending on the specific implemented model, the anisotropic compartments were optionally convolved with a Watson or a Bingham distribution to account for fiber orientation dispersion. A summary of the multicompartment models that were submitted and the components they are based on is shown in Table 4.

**Table 4:**
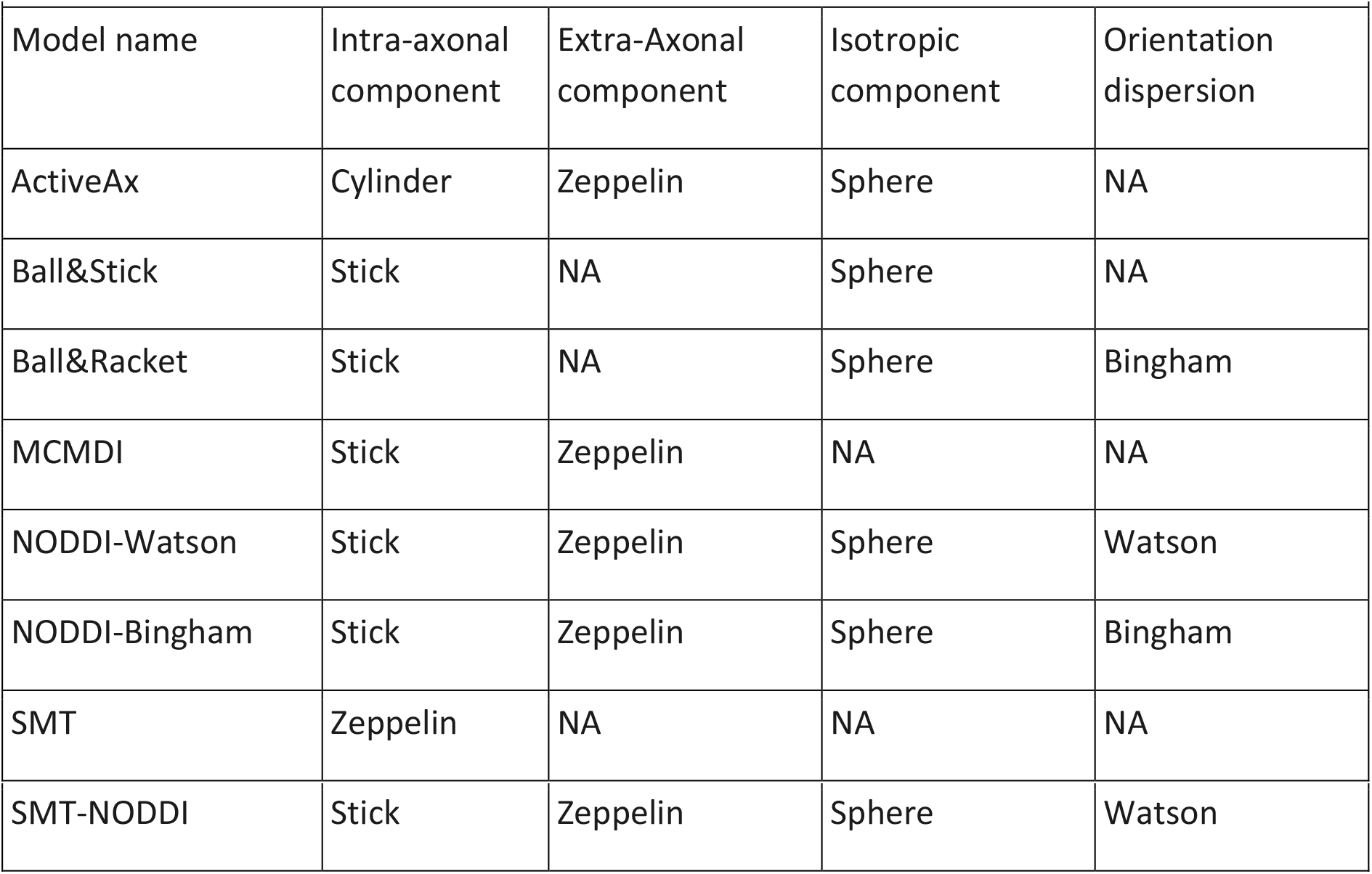
An overview of the diffusion models used to represent the individual components of the considered multicompartment models.

For all the above mentioned models, the parallel diffusivity of the anisotropic compartments was set to 1.7×10^-3^mm^2^/s, whereas the diffusivity of the isotropic compartment was set to 3×10^-3^mm^2^/s, as previously suggested (Zhang et al. 2012; Kaden et al. 2016; Behrens et al. 2003; Sotiropoulos et al. 2012). The perpendicular diffusivity of the anisotropic compartments was linked to the parallel diffusivity via the tortuosity constraint (Szafer, Zhong, and Gore 1995; Zhang et al. 2012).

#### Parametric representations

This family of methods focuses on expressing the dMRI signal as a function of mathematical signal basis without biophysical hypotheses. A popular signal representation is the simple harmonic oscillator reconstruction (SHORE)(Ozarslan et al. 2009). The SHORE basis has some degrees of freedom such as the order and a scaling factor. The SHORE method was applied to predict all the 4 provided data types (SDE-MS, SDE-GRID, DDE, DODE) in combination with a BFGS fit (Nath, Lyu, et al. 2019) that can be utilized to achieve the best fit of the scaling parameter. The same method was used for all 4 types of data with harmonics of orders 6, 8 and 12.

MAP-MRI is a signal representation-based technique that expresses the diffusion signal in q-space and parametrizes its Fourier transform –the mean apparent propagator (MAP)– in terms of a series involving products of three Hermite functions, thus generalizing the 1D-SHORE technique to three dimensions. Unlike in 3D-SHORE, an anisotropic scaling parameter is employed in MAP-MRI, making it an extension of DTI for representing the signal at large q-values. In this challenge, we received submissions from two different teams based on a Laplacian regularized version of MAP-MRI. Both submissions (MAP-MRI+Reg) predicted the unseen data points using penalized least squares and SHORE basis of order 8.

#### Neural networks

Convolutional neural networks (NeuralNet) are increasingly being used for tasks such as quantification and signal representation. In this case, the output of the networks corresponded to the dMRI signal.

The first family of submissions that we received is based on feed-forward networks with a single hidden layer of 50 neurons and sigmoid activation functions. These networks were trained on 80% of the measurements and validated on 20% by minimizing the mean squared error of the predictions with the *AdamW* algorithm. For the learning phase, a learning rate of 0.005 and 20000 epochs were used. To predict the SDE signals, the normalized components of the gradient (3 values) and the b-value were provided as inputs. For the DDE and DODE acquisitions, the gradient strength, the normalized components of the two gradients (6 values), the b-value, and the components of the b-matrix (6 values) were concatenated into one input vector of length 14. The training and prediction phase were repeated independently for each of the individual signal and data types.

The second family of submissions we received was based on neural networks with reinforcement learning (NN+Reinf) (Zoph and Le 2016; Williams 1992). A neural architecture search (NAS) was implemented to search the optimal 7-layer feed-forward model with ReLU activations for dMRI signal prediction given the acquisition parameters. The search space of NAS is the number of nodes in each of the seven layers in the set [8, 16, 32, 64, and 128]. and fit the training data better. And the number of neurons in each of the seven layers belongs to [8, 16, 32, 64, and 128]. For training, the initial learning rate was set to 0.01, and the adam optimizer was used.

## Data analysis

The data analyses were performed separately for SDE-MS, SDE-GRID, DDE and DODE. For each encoding type and signal, we evaluated the min-max interval of all predictions, and their 25th to 75th percentile confidence interval. For each signal, we determined the best prediction as the one achieving the lowest mean squared residuals (MSE), and visually investigated the residuals. As the MSE only represents one of the possible metrics that can capture the goodness of the signal prediction, we also determined the variance of the residuals and the bayesian information criteria (BIC) associated with the prediction of each individual signal. These results can be found in the supplementary material Table S2 and S3.

Subsequently, the distribution of the residuals of each model were investigated by means of boxplots. At this stage, the residuals from all the 5 voxels were considered together. A ranking of all considered models was derived according to increasing MSE, then the best prediction model was established per encoding type. The residuals of the best prediction model were investigated as a function of the b-value, diffusion time (DDE only) and encoding frequency (DODE only), to understand whether all diffusion encodings were predicted with comparable accuracy and precision. In the subsequent analysis, the 5 voxels were studied separately. Specifically for the SDE-MS data, an additional analysis was performed to evaluate how the best model predicted the directional dependent information of the shell acquired at b = 4000s/mm^2^. To this end, the measured data and the best signal prediction were projected on the unit sphere using spherical harmonics of order 12, then the prediction error was evaluated. The diffusion tensor imaging model was included in this step for reference.

## Results

### Signal representation of SDE-MS data

The geometric average of the SDE-MS signals and an overview of the predictions is shown in Figure 2. In general, both the average and the best fitting method predicted the average signal decay without apparent biases, with the exception of the average fit of the low diffusion weightings of signals 3 and 5.

**Figure 2.**
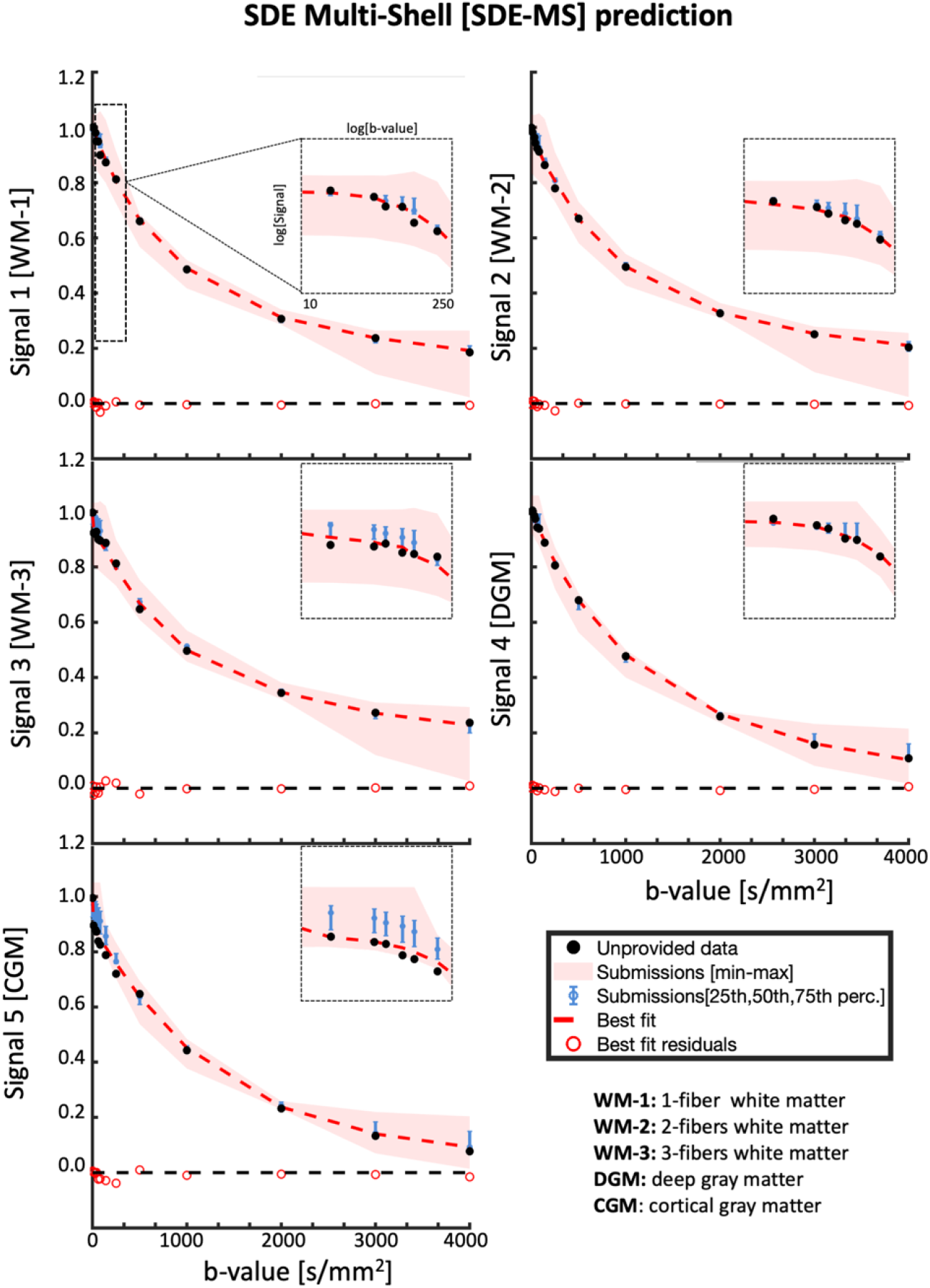
Signal decay as a function of the b-value of the averaged SDE-MS data over different directions, for the unprovided measurements. The black dots represent the unprovided data, the red shaded area represents the min-max of the submissions, the blue error bars represent the 25-75 percentile, the solid red curve represents the best fit and the red circles represent the residuals of the best fitting model. The 5 different plots illustrate the predictions of the different signals.

On average, the confidence interval of the submissions (blue error bar) is centered on the geometric average of the data for diffusion-weightings 400 < b < 4000 s/mm^2^ for all 5 signals. The prediction of data at b = 4000 s/mm^2^ (which was not provided in the training data) was overall accurate in WM (signals 1-3), but showed a small and consistent over-estimation in DGM and GM. The signal measured at low diffusion weightings (i.e. b < 200 s/mm^2^) was on average accurately predicted in the WM voxel containing up to 2 crossing fibers and in deep gray matter, but not in the complex WM-configuration (Signal 3) and in cortical GM (Signal 5). The min-max range of the predictions highlights that data at b = 2000 s/mm^2^ is predicted on average with the lowest uncertainty, whereas the largest spread is observed at b < 200 s/mm^2^ and b = 4000 s/mm^2^. The best predicting models for signals 1-5 are SHORE, MAP-MRI+Reg, MAP-MRI+Reg, Ball&Racket and NeuralNet, respectively.

Figure 3 shows the average residuals of all tested signal predictions when considering the 5 provided signals together. The predictions of the first 7 methods were remarkably accurate, as shown by the tight confidence interval of the residuals of about 0.03 of the measured data, which occasionally reached values up to 0.1. MAP-MRI provided the best overall signal prediction with average MSE 0.00236 ± 0.00035 (Supplementary Material Table S1). When comparing the residuals of this prediction to those from the remaining models, we found that only NeuralNet provided equally distributed residuals (sign test, p<0.05). When looking at the average residuals of each signal individually (colored dots in the boxplots of Figure 3), a tendency towards a better prediction of WM signals as compared to CGM and DGM was observed.

**Figure 3:**
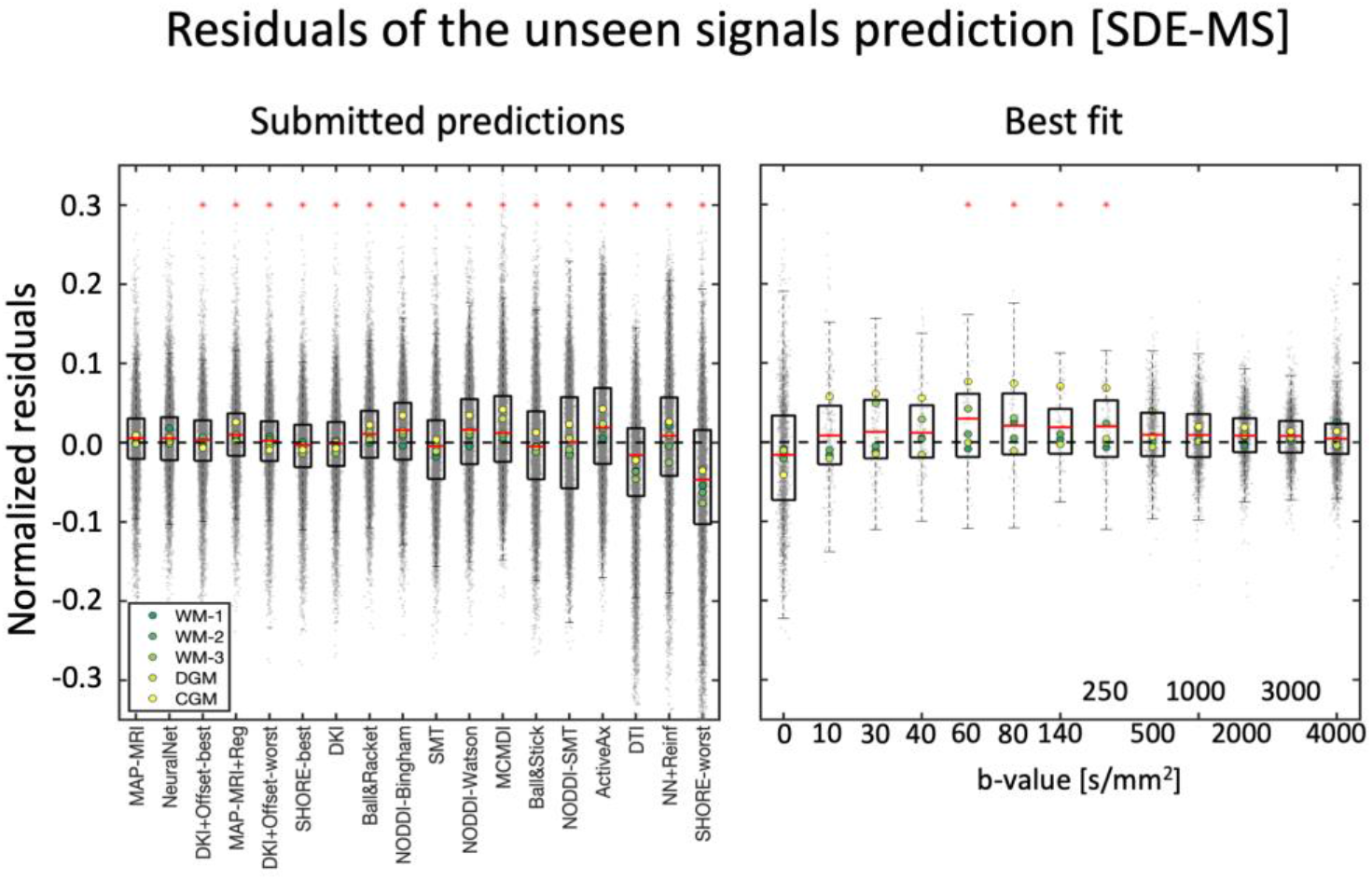
Left) Boxplots of the normalized residuals (gray dots) of each prediction of SDE-MS data, when pooling together all 5 signals. Right) The normalized residuals of the best prediction (MAP-MRI) over individual diffusion weightings. The red asterisks on the left panel indicate predictions significantly different from the best prediction, whereas those on the right indicate that residuals at a specific diffusion weighting show a significantly non-zero mean.

To understand how the fit of the best prediction method (MAP-MRI) varied as a function of the diffusion-weighting, we evaluated the value of its average residuals for all 5 signals together for each shell independently. In general, the average residuals of most shells were close to zero, but a significant overestimation for b-shells 60 <= b <= 250 s/mm^2^ was observed (one sample t-test, p < 0.05). The average prediction error was on average less than 0.05 for all shells, but errors up to about 0.2 can be observed for the non-weighted data and the unprovided shell at b = 4000 s/mm^2^.

Having established that model MAP-MRI provided the best average fit, we set to investigate how this method could predict the angular information of the unprovided shell at b = 4000 s/mm^2^, and included a prediction with DTI and the average prediction of all methods for reference. This angular resolution analysis is shown in Figure 4.

**Figure 4:**
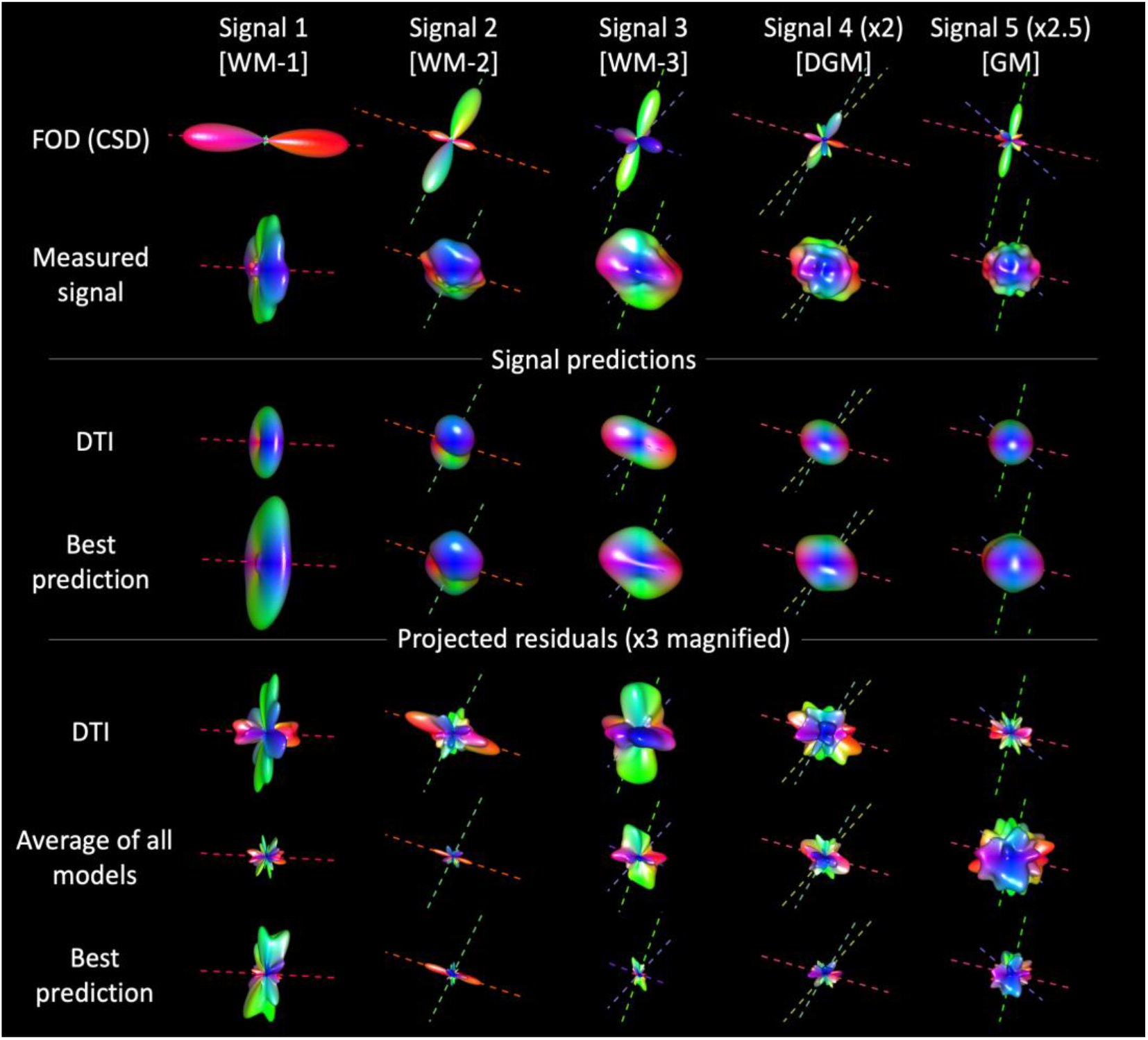
3D visualisation of the fiber orientation distribution (FOD), a projection on the unit sphere of the measured signal, of the signal predictions with DTI and with the best prediction model as well as of the residuals determined with DTI, average of all models and with the overall best predicting model for the unprovided SDE-MS data at b = 4000 s/mm^2^

All methods could well-predict on average the donut-shaped 3D representation of WM-1, and the average residuals of this signal are smaller than those of the best fitting method. The DTI prediction of WM-1 well captured the overall shape of the signal, but also showed large errors in specific directions, likely due to unaccounted signal restrictions at this diffusion weighting. The best prediction was much better than the average prediction as well as the DTI one in the more complex configurations WM-2 and WM-3. In the WM-signals, the largest angular errors were observed in directions parallel and perpendicular to the main diffusion directions derived with CSD, as expected. In DGM and CGM, all methods performed overall equally well and the residuals had an almost isotropic distribution.

### Signal representation of SDE-GRID data

Figure 5 reports the average signal predictions for SDE-GRID after binning closely spaced diffusion-weightings to enhance clarity. In general, the SDE-GRID was well predicted by most submissions, as highlighted by the tight min-max and confidence intervals. Larger prediction variance can be observed at low (b < 1000 s/mm2) and large diffusion weightings (b > 6000 s/mm2) than in the intermediate range. The best predictions of signals 1 and 2 were achieved with DKI+Offset, whereas MAP-MRI+Reg provided the best prediction of signals 3-5 and the best overall prediction with MSE 0.00260 ± 0.00043 (Supplementary Table S1).

**Figure 5.**
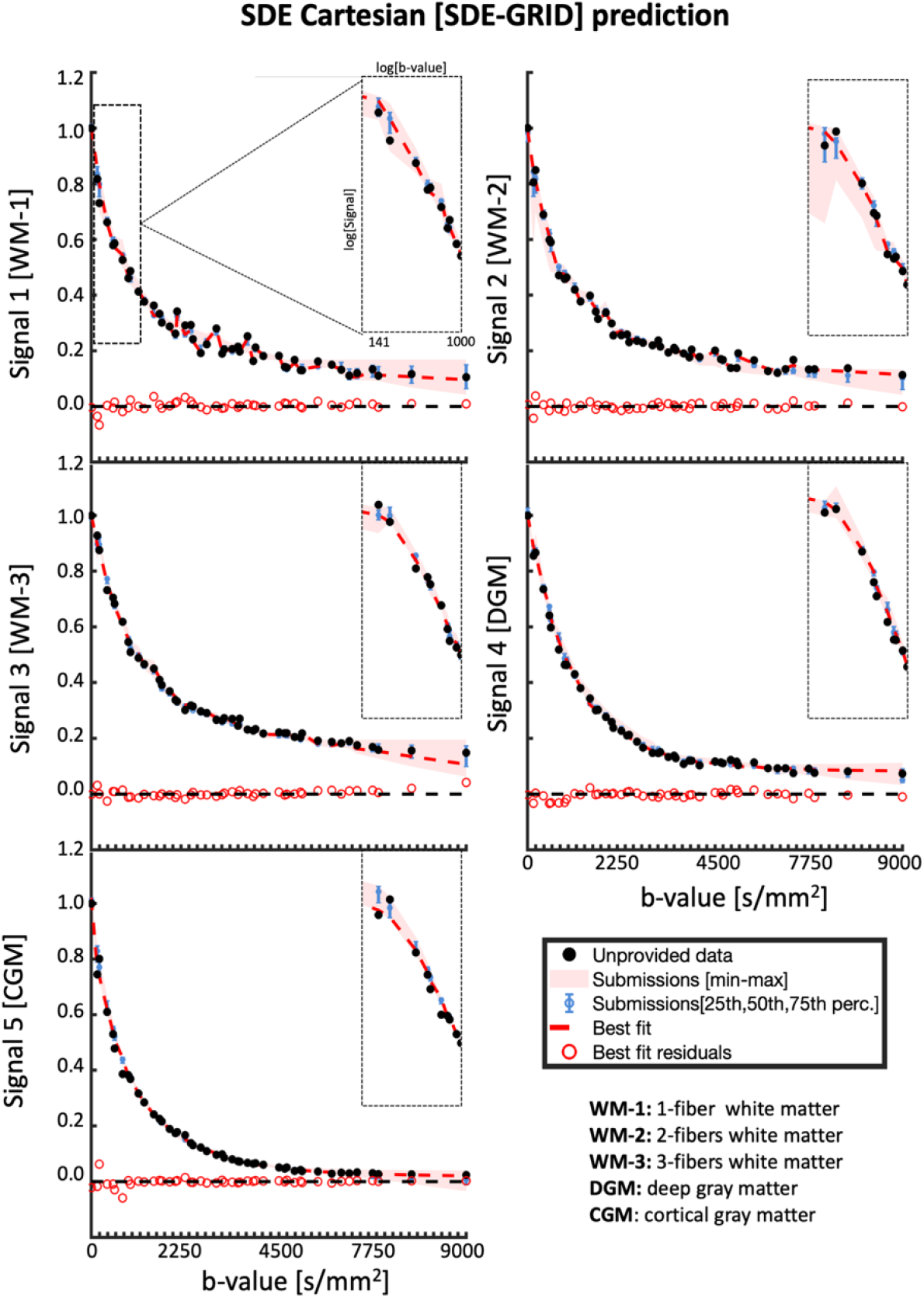
Signal decay as a function of b-value of the averaged SDE-GRID data over different directions, for the unprovided measurements. The diffusion weightings were rounded to the closest multiple of 100 before averaging to enhance clarity. The black dots represent the unprovided data, the red shaded area represents the min-max of the submissions, the blue error bars represent the 25-75 percentile, the solid red curve represents the best fit and the red circles represent the residuals of the best fitting model. The 5 different plots illustrate the fits of the different signals.

The boxplots of residuals of the SDE-GRID predictions ranked by MSE are reported in Figure 6. The first 7 submissions predicted the signals accurately, without visible biases both at average level as well as in specific tissue-types, and most prediction errors were in the range of ±0.03 with values occasionally up to 0.1, similar to what was previously observed for SDE-MS. When considering all predictions together, MAP-MRI+Reg provided the best overall prediction, but predictions with DKI+Offset and NeuralNet can be considered comparable according to a signed rank test. When analyzing the average prediction residuals of MAP-MRI+Reg for the binned diffusion weightings, it is appreciable that most data was well predicted without biases and errors below 0.05, with the exception of b <= 800 s/mm^2^, b around 3000 s/mm^2^ and b > 6000 s/mm^2^.

**Figure 6:**
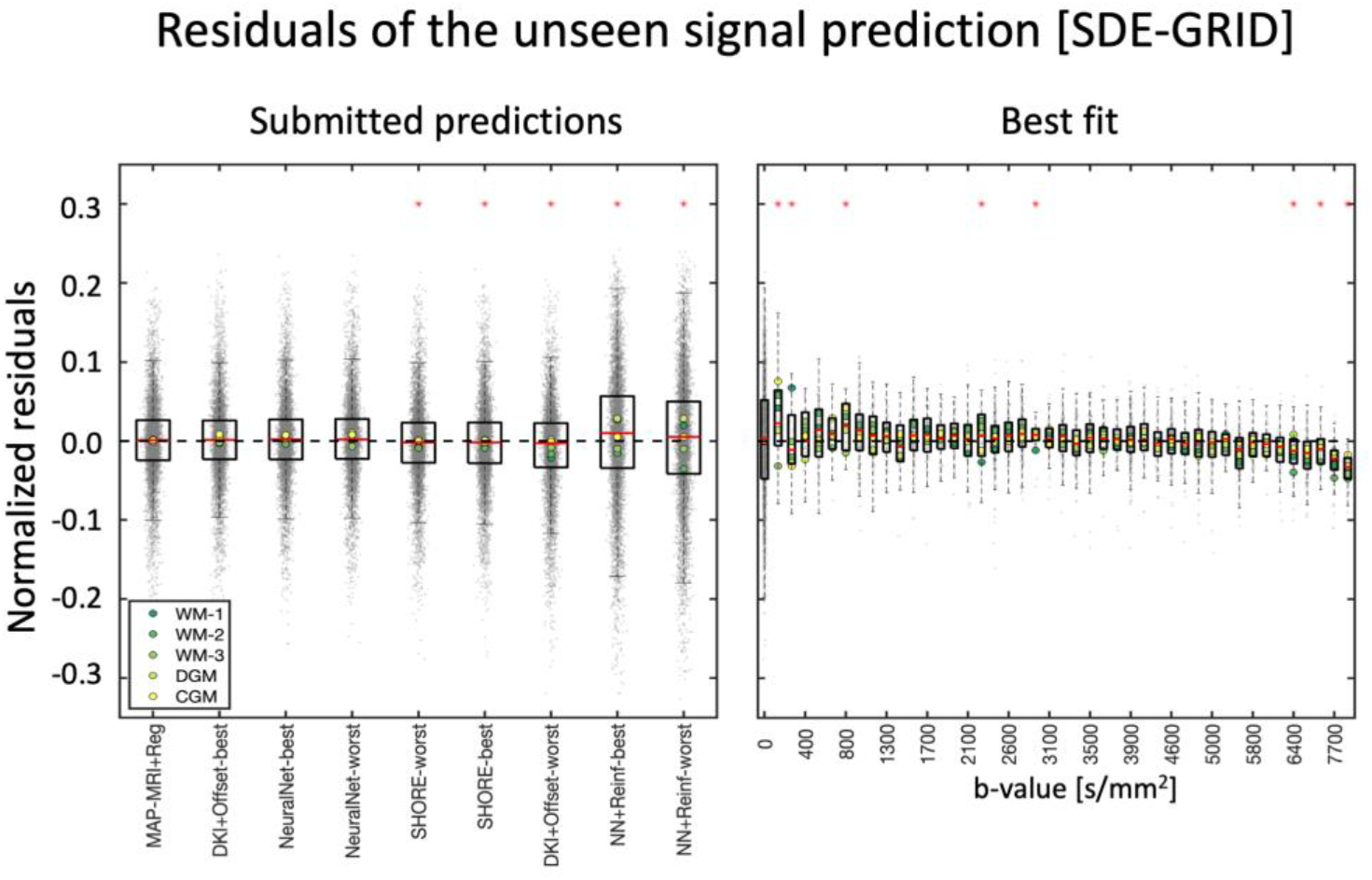
Left) boxplots of the normalized residuals (gray dots) of each prediction of SDE-MS data, when pooling together all 5 signals. Right) The normalized residuals of the best prediction (MAP-MRI+Reg) over individual diffusion weightings. The red asterisks on the left panel indicate predictions significantly different from the best prediction, whereas those on the right indicate that residuals at a specific diffusion weighting show a significantly non-zero mean.

### Signal representation of DDE and DODE data

Figure 7 shows the best and average signal predictions of the unseen DDE and DODE data for the 5 different voxels. Figure 8 presents the normalized residuals for the different submissions, averaged over voxels, b-values and diffusion times/frequencies while Figure 9 shows the normalized residuals for the best fitting model as a function of the b-value.

**Figure 7.**
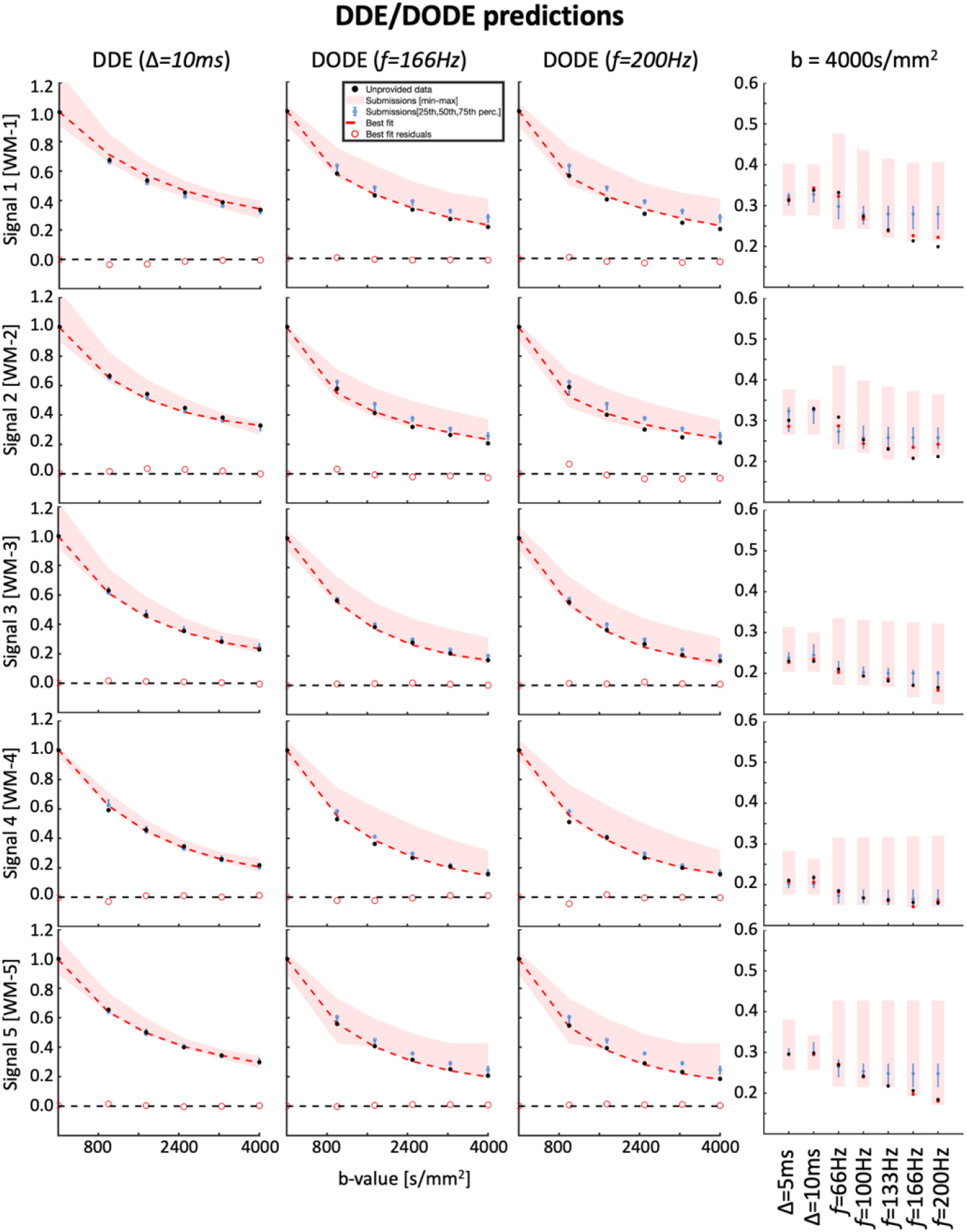
The first three columns show the signal decay as a function of b-value of the geometrically averaged DDE and DODE data over different directions, for the unprovided measurements. The fourth column shows the geometric average of the signal measured at b = 4000 s/mm^2^ for different diffusion times Δ (DDE) and oscillation frequencies *f* (DODE). The black dots represent the unprovided data, the red shaded area represents the min-max of the submissions, the blue error bars represent the 25-75 percentile, the solid red curve represents the best fit and the red circles represent the residuals of the best fitting model. The 5 different plots illustrate the fits of the different signals.

**Figure 8:**
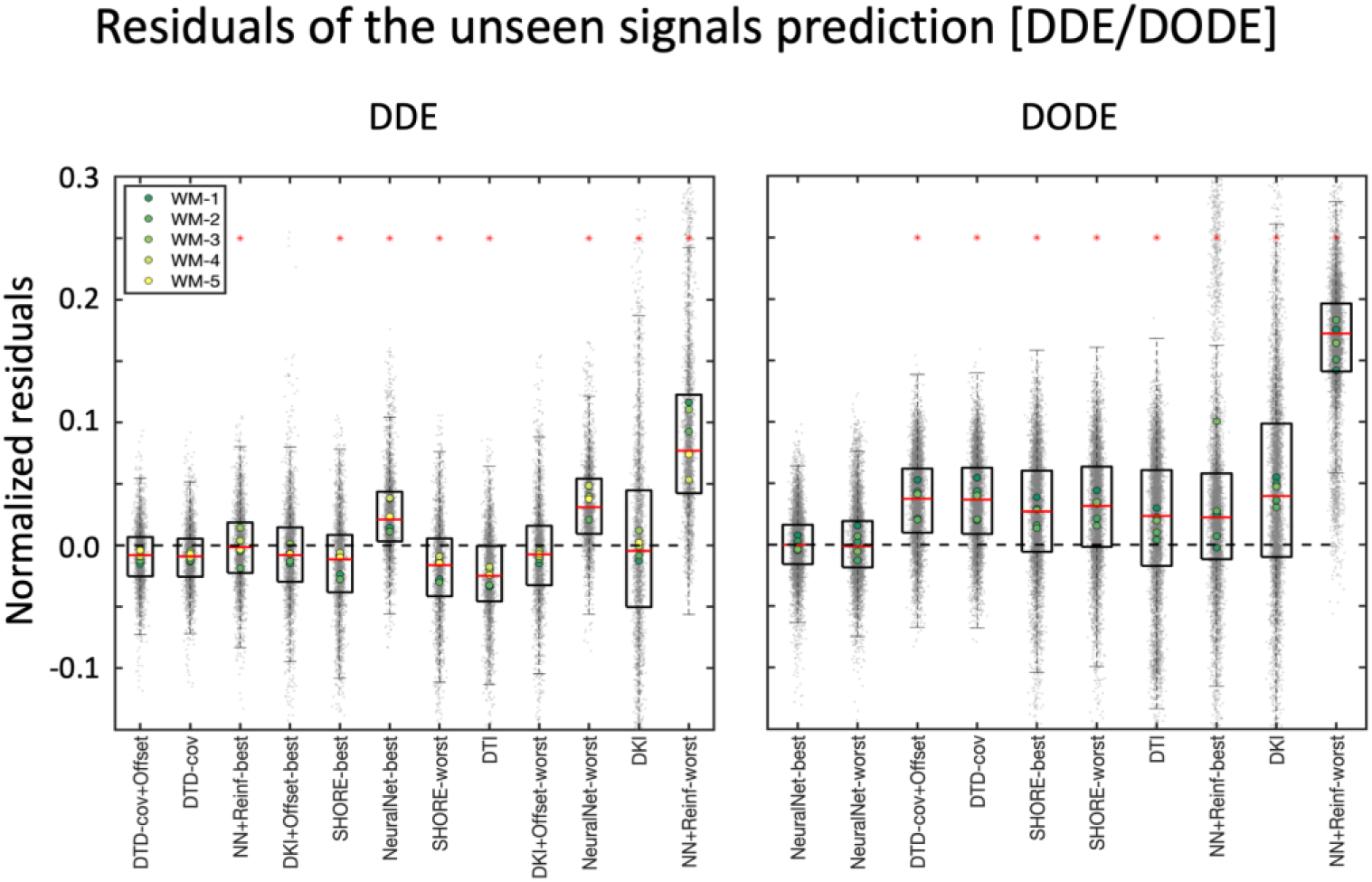
The boxplots of the normalized residuals (gray dots) of the DDE (left) and DODE (right) predictions. The red asterisks on the panels indicate predictions significantly different from the best prediction. The first 5 DDE predictions perform reasonably well as shown by the value of most residuals being well-below 0.1, although a trend towards the overestimation of the signal could generally be observed.

**Figure 9 -.**
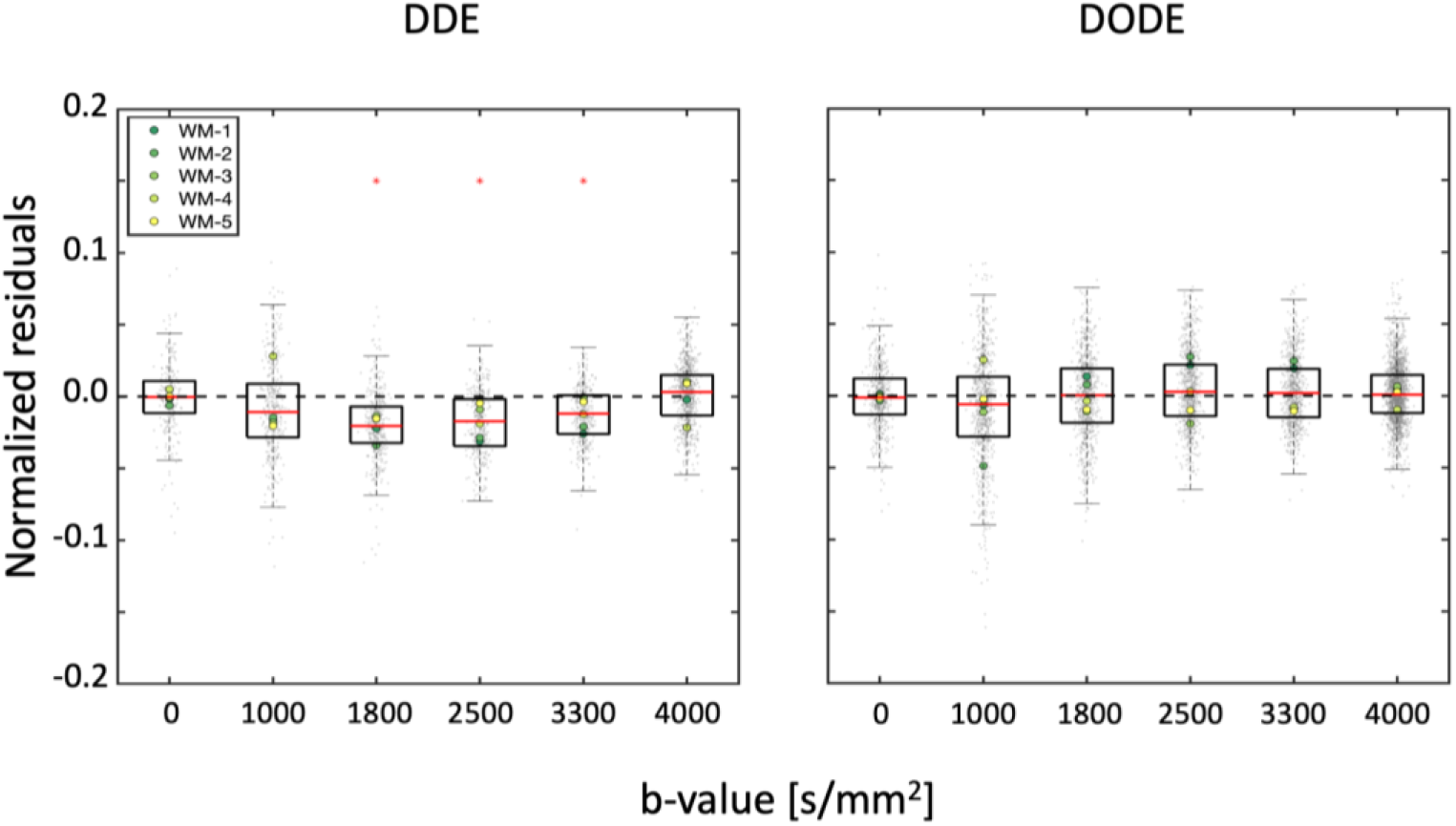
Right) The normalized residuals of the best prediction of DDE (DTD-cov+Offset) and DODE (NeuralNet-best) over individual diffusion weightings. The red asterisks indicate that residuals at a specific diffusion weighting show a significantly non-zero mean. With DDE data, this was observed only for b-values between 1800 and 3300 s/mm^2^, whereas no biases were observed for DODE.

For DDE we see that the prediction of the directionally averaged signal is well aligned with the measured data with the DTD-cov+Offset providing the best prediction with MSE 0.00072 ± 0.00023 (Supplementary Table S1). Nevertheless, other methods such as DKI, SHORE and neural networks also performed reasonably well in providing unbiased predictions, but with visibly larger errors. In general, we see that the prediction of the higher b-values (> 2500 s/mm^2^) is better than the prediction of the lower b-values (< 2500 s/mm^2^). For DODE data, the best prediction comes from NeuralNet-best with MSE 0.00070 ± 0.00036, whereas the majority of the submissions overestimate the signal, especially for b-values larger than 1750 s/mm^2^. For both DDE and DODE, the predictions show similar trends in the 5 different white matter voxels. For the DODE data, both frequencies also show similar trends of the predicted signal.

## Discussion

We have evaluated the generalizability of existing dMRI methods at predicting diffusion-weighted data measured with SDE-MS, SDE-GRID, DDE and DODE by analyzing 80 submissions to the MEMENTO challenge from 7 different teams. In general, our analysis suggests that models predicting SDE-MS and SDE-GRID data generalized the easiest to unseen diffusion encodings, whereas the prediction of DDE and DODE seems more challenging. Within the domain of SDE, the worst prediction was observed in correspondence of low and very strong diffusion weightings.

### Trends in SDE data predictions

The large majority of the analyzed submissions predicted SDE data, with a considerable preference for SDE-MS over SDE-GRID, which also reflects the overall larger number of studies which employ shell data. When looking at SDE-MS, we can observe that the majority of submissions could well predict the global signal decay, and 14 out of 18 predictions had a median error smaller than 0.04. Of these, however, only 7 had an interquartile range (25th-75th percentile) of the residuals smaller than 0.05 in absolute value, which suggests how the prediction of the isotropic component of the signal decay (which captures the average decay of a given diffusion weighting) is an easier task than the prediction of the anisotropic component. Interestingly, the 7 predictions with the lowest MSE can account for complex fiber configurations such as 2+ crossing fibers, whereas predictions with single-fiber based methods result in higher MSE. Looking at the angular analysis reported in Figure 4, it becomes clear that MAP-MRI provides the best prediction of both SDE-MS and SDE-GRID by well representing the signal in voxels with complex fiber configurations (WM-2, WM-3) as well as in DGM, whereas the prediction error in voxels with a single fiber population (WM-1) or almost isotropic diffusion (CGM) is worse than the average submission.

A second aspect regarding the analysis of SDE data is the dependence of the prediction accuracy and precision on the diffusion weighting and on the specific tissue type. Our results suggest that current dMRI methods can well represent and predict dMRI data with commonly used diffusion weightings. Indeed, we observed that most submissions could predict SDE data remarkably well within the range of commonly employed diffusion weightings (e.g., 1000 <= b <= 4000 s/mm^2^), whereas the prediction of low (b < 800 s/mm2) and strong (b > 6000 s/mm^2^) diffusion weightings was generally less accurate. While the latter might originate from Rician-related biases, it might also highlight the limited specificity of existing models to genuine components such as perfusion contributions at low diffusion weightings (Le Bihan et al. 1988; Pasternak et al. 2009) and WM-restriction at strong diffusion weightings (Cohen and Assaf 2002). In this context, a trend towards a worse prediction of the DGM and CGM signals as compared to WM signals emerges with SDE-MS and, to a lesser extent, with SDE-GRID. This seems to be mostly driven by the inaccurate prediction of the signal measured at low diffusion weightings where the sensitivity to blood pseudo-diffusion is maximal, which once again suggests a lack specificity at taking into account specific properties of the GM like its higher perfusion as compared to WM (Ahlgren et al. 2016). This holds also for the best prediction (MAP-MRI) as shown by the significant overestimation of the signal at b < 250s/mm^2^, where the contribution of pseudo-diffusion effects becomes non-negligible. A bias in the prediction of SDE-GRID with MAP-MRI is also revealed for b > 6000 s/mm^2^. While this effect might be partially explained by Rician noise, the observation that its effect is larger in WM than GM, and that it grows in magnitude with fiber complexity being the largest for WM-3, suggests the presence of a genuine unaccounted trend in the signal. Interestingly, smaller errors are observed for the prediction of SDE-GRID than of SDE-MS on average, and even methods providing visibly biased SDE-MS predictions such as SHORE-worst, performed well at predicting SDE-GRID. This might be explained by the larger range of unique diffusion weightings included in SDE-GRID providing less redundant information than many measurements in few shells, or to the larger minimum diffusion weighting included in SDE-GRID (b = 141 s/mm^2^) as compared to SDE-MS (b = 10 s/mm^2^).

### Trends in DDE and DODE data predictions

The prediction of DDE and DODE data seems more challenging than that of SDE. Indeed, the signal measured with DDE and DODE is encoded with additional dimensions as compared to SDE, namely parallel and orthogonal gradient orientations within one measurement, leading to linear and planar b-tensors, respectively, as well as different diffusion times and oscillation frequencies. In the challenge design, this aspect was stressed by requiring the prediction of unseen diffusion-weightings and gradient directions for 1 completely unseen diffusion time (DDE) and 2 unseen oscillation frequencies (DODE) provided a training set encoded with a different diffusion time and 3 different oscillation frequencies, which is arguably a harder task than the prediction of SDE data. The first take home message from the analysis of the DDE and DODE predictions is the need to take into account the additional encoding dimension, which is in line with our expectations given that the previous analysis of the data showed a clear diffusion time/frequency dependence over this parameter range (e.g. Fig. 9 in (Ianus et al. 2018)). For models which do not account for diffusion time / frequency (e.g. DTI / DKI / DTD, etc), we expect the predictions for the unseen DDE with Δ = 10 ms to be the same as the model prediction for the provided DDE data with Δ = 5 ms. Since for DDE only one diffusion time was provided, the neural networks could not learn the effect of this parameter, so in practice none of the submissions could account for diffusion time. For the unseen DDE data with Δ = 10 ms, the best model from the current pool of submissions is DTD-cov+Offset, which nevertheless resulted in a small bias between the measurements and the predictions. The prediction errors of the directionally averaged signal provided by the best model are larger for intermediate b-values (e.g., b = 1800 – 3300 s/mm^2^) than for the highest b-value of 4000 s/mm^2^. This might be the case because the effect of diffusion time on the signal over this parameter range becomes smaller at higher b-value, for instance due to a smaller contribution of extracellular space.

For DODE data, multiple frequencies were provided for training. The best DODE predictions were achieved with methods able to account for the frequency implicitly, such as in the case of neural networks, whereas most submissions overestimated the measured signal. Interestingly, the error increases on average with the oscillation frequency, and even the best prediction, NeuralNet-best, could not predict well the data at b = 4000 s/mm^2^ for *f*=200Hz. All submissions but one achieved a 25th to 75th percentile error below 0.1, but 8 out of 10 submissions exhibited a consistent median error of about 0.04. Altogether, this suggests in our opinion that we currently do not fully understand how to properly model the effect of frequency, and highlights the need for further research in the optimal modelling of DODE data.

### Deep learning-based methods

The application of machine learning and deep learning is currently booming across all science fields dealing with large data. MRI and dMRI are no exception, and in this very own challenge we have received 32 submissions based on deep learning methods next to established signal representations and biophysical models, which represents 40% of the total submissions. Interestingly, neural network-based methods provided accurate predictions for different diffusion encodings, achieving the 2nd best prediction for SDE-MS, the 3rd best prediction for SDE-GRID and DDE, and the best prediction of DODE data. The latter performance is remarkable, because NeuralNet-best and NeuralNet-worst provided the only two unbiased DODE predictions, showing ability to learn the relation between the diffusion signal and its encoding parameters, including the oscillation frequency, without the need for explicit modelling. These results certainly showcase the potential of these methods, and support their applicability as excellent interpolators, able to learn data features from a rich dataset and to provide good predictions of unseen data - within the boundaries of their training set. Most deep-learning based methods do not quantify metrics that can be used to extract properties of in-vivo tissues, and are thus unlikely to spread into clinical use at the current moment. Nevertheless, their good prediction performance make them favorable for tasks where a direct manipulation of the signal is required, such as denoising, artefact removal or even data augmentation, and their application in combination with classic methods might prove advantageous to enhance the quality of the results or to shorten acquisition time.

### Importance of hyperparameters and user choices

Our analysis highlights how user choices and hyperparameters can remarkably affect the prediction accuracy. The SHORE method, for example, achieved both one of the best predictions of SDE-MS data as well as the worst, with an average error difference between the two of about 8%. Similarly, the addition of a degree of freedom to DKI or DTD-cov (i.e., +Offset) appreciably improved its accuracy. DKI+Offset, for example, predicted the SDE-MS and DDE data with an average prediction error 10% and 92% smaller than DKI, respectively. Large variability in the prediction performance was also observed for neural networks methods, which flexibility in design allows the implementation of very diverse architectures with performance strongly influenced by the optimization of hyperparameters. Altogether, we believe that this highlights the importance of not only reporting the specific method used for data analysis, but also to explain the choice of its hyperparameters and, where possible, to share its implementation to maximize the comparability of results obtained in different studies. Next to user settings and hyperparameters, model assumptions are an important factor that might limit the generalizability of a prediction method. This is especially the case for multi-compartment models based on physiological assumptions, including fixing the intra-cellular diffusivity to literature values or using tortuosity principles to constraint the cellular fraction, as recently suggested (Lampinen et al. 2019; Henriques et al. 2019).

### Link to previous challenges

Various dMRI challenges have been organized in the past (Ferizi et al. 2017; Schilling et al. 2019; Pizzolato et al. 2020), with a focus on different data aspects. For instance the ones described by Schilling et al. 2019 focused on tractography, while others have focused on data prediction, including diffusion measurements at multiple echo times (Ferizi et al. 2017) as well as inversion times (Pizzolato et al. 2020). In terms of challenge requirements, the one organized by (Ferizi et al. 2017) is the most similar to the current one, as it also evaluated the submissions based on the prediction of unseen SDE data shells.

Nevertheless, the majority of models submitted to the SDE part of this challenge do not overlap with the models included in the previous challenge, making a direct comparison difficult. One notable exception is MAP-MRI, which performed very well for the current SDE datasets, but not for the one considered by (Ferizi et al. 2017) which included multiple diffusion and echo times. In that case, models which accounted for multiple compartments with different T2 properties provided better estimates, an effect not considered here. In addition to the development of more models and signal representations, the current work investigates generalizability over a wider range of diffusion weightings, angular directions, and diffusion times.

### Limitations

Some limitations of this study should be acknowledged for a more comprehensive interpretation of the presented results.

#### Data limitations

Firstly, the SDE-MS/SDE-GRID and DDE/DODE data have been acquired in very different settings. The SDE data were acquired as part of the MASSIVE data (Froeling et al. 2017b) and represent an unique collection of thousands of unique diffusion measurements in an in-vivo human brain at 3T, but were spreaded in 18 acquisition sessions - which might introduce additional variability in the data - and are characterized by an overall modest SNR at b = 0s/mm^2^ (~15). Conversely, the DDE/DODE data were acquired in an ex-vivo mouse brain with a state-of-the-art 16.4T scanner, and characterized by very high SNR. Consequently, the generalizability of our results to DDE/DODE data acquired in in-vivo humans at 3T requires thus further research. While we expect our findings to generalize to individuals with similar characteristics, e.g., healthy adult humans (SDE) and mice (DDE/DODE), some results might be driven by the unique characteristics of these brains, and will likely not extrapolate well in presence of pathology, as well as in infants and elderlies.

#### Data selection

A further element of variability in the comparison of the two datasets is introduced by the different criteria used for the selection of the signals: on the SDE data, we sampled signals from WM voxels with different configurations (1 to 3 fibers), but also included GM voxels, which allowed us to investigate tissue-specific performance. As a consequence of this choice, any submissions regarding SDE-MS and SDE-GRID needed to perform well in both WM and GM to achieve a good score. Differently, all signals sampled from the DDE/DODE datasets were located in WM and offer a more thorough overview of the prediction performances across different fiber configurations - including regions with known fiber fanning - but no insights into their applicability to GM. For all of the above, the prediction performance obtained by the submissions on SDE and DDE/DODE should not be directly compared. In the selection of the voxels, we attempted to avoid tissue interfaces in order to minimize partial volume effects between different tissue types, but these cannot be ruled out and are expected to be more detrimental for methods not explicitly dealing with partial volume effects. Nevertheless, we argue that some extent of partial volume is ubiquitous in brain dMRI applications, and that taking it into account is likely part of the challenge to accurately model the dMRI signal.

#### Challenge evaluation

As a community challenge, we chose to calculate a single metric (the mean squared error) in order to determine a “winning” algorithm. Other choices of the score criteria were possible, and would likely result in a different ranking. For example, according to modelling theory it would seem more appropriate to investigate a goodness of fit criteria as the Bayesian information criteria rather than considering the signal residuals alone, to penalize signal overfitting (Supplementary Material Table S2 and S3). However, it is arguable that these kinds of metrics are not suitable to characterize methods based on machine learning / deep learning where thousands to millions of parameters are fitted, and that the mean squared error captures, in its simplicity and limitation, the basic ability to predict an unseen signal. Nevertheless, doing well in the current challenge does not automatically guarantee that these algorithms are the most appropriate models in all cases. Here, we have focused on the ability to explain (i.e. predict) the signal over a wide range of diffusion weightings, diffusion times, and frequencies. Furthermore, some modelling approaches in this study may be suitable only for a subset of the wide range of acquisitions in this database, and may be more/less sensitive at different areas of the diffusion sensitization space. Tensor-based models such as DTI and DKI, for example, are known to well fit data in the range b = 800-1200 s/mm^2^ and b = 1000-3000 s/mm^2^, respectively. Unsurprisingly, in this challenge we indeed observe very large residuals for the DTI model at b < 500 s/mm^2^ and b > 2000 s/mm^2^, and for the DKI model at b < 800 s/mm^2^ and b > 5800 s/mm^2^, respectively, which penalize the final scoring of these methods. Another aspect that might influence the evaluation is the pre-processing of the data, which is well-established to have a major impact on the subsequent data analysis. To rule out its potential confounding effect on our results, we have provided the participants with standardized - already pre-processed data, but the inclusion of additional pre-processing steps (i.e., denoising, Gibbs ringing correction, outliers replacement, etc.) might have resulted in different prediction performance and “winners”.

#### Lack of validation

The ability of a method to accurately predict unseen data offers a measure of fidelity to the underlying tissue microstructure, but it is by no means a substitute for validation efforts that compare signal models to the actual biological tissue structure obtained through orthogonal measurements such as, for example, high-resolution microscopy. Thus, we argue that the appropriateness and specificity of the tested methods cannot be adequately captured by signal fitting alone, and requires external validation, which is particularly critical in the case of biophysical multicompartment models. In addition to empirically assessing data, future work should continually strive to validate these measures against orthogonal information through simulations, physical phantoms, and animal models of tissue microstructure in order to paint the complete picture of the models successes and abilities.

### What is a good model?

As described in past challenges (Panagiotaki et al. 2012; U. Ferizi et al. 2015; Ileana O. Jelescu et al. 2020) and in reviews (Ileana O. Jelescu et al. 2020; Dmitry S. Novikov, Kiselev, and Jespersen 2018), a good model or signal representation must well-capture trends in the signal (explain seen signal and predict unseen signal), and also have stability and robustness of fit (Ileana O. Jelescu et al. 2016), *for the appropriate signal regime*.

On the other hand, a good model fit to the data, and ability to predict unseen data, does not guarantee that the estimated model parameters have a sensible physiological meaning. Similarly, a visually appealing map of quantitative indices also does not equate to a “good model”. While the “best” model is the one that well-explains the underlying physiology that the signal is sensitive to (within the experimental design), the process of converging on the most-appropriate model is complex, and examining the generalizability of the model to various diffusion sensitizations is only one step in that process. This specific step lends insight into the information uncovered and captured in the signal, and successes and limitations of various attempts to describe the signal.

### Conclusions

We have reported the results of a community effort to investigate the generalizability of existing methods at predicting unseen diffusion MRI signals collected over a large range of diffusion encodings. Our results highlight that existing models perform well at predicting SDE data in white matter and, to a lesser extent, in grey matter. Conversely, future work is needed to better understand and model the information content of DDE and DODE data. Next to the method choice, hyperparameters play a key role in the generalizability of fit methods, highlighting the importance of their optimization, and of reporting their values to support reproducibility. These challenge results serve not only as a snapshot of the current status quo in the field, but also as an openly available benchmark to support the development of novel methods.

## Acknowledgements

The work by Inria co-authors was partially funded by the European Research Council (ERC) under the European Union’s Horizon 2020 research and innovation program (ERC Advanced Grant agreement No 694665: CoBCoM - Computational Brain Connectivity Mapping, P.I. Rachi Deriche) and by the French government, through the 3IA Côte D’Azur Investments in the Future project managed by the National Research Agency (ANR) with the reference number ANR-19-P3IA-0002.

Marco Palombo, Daniel C Alexander and Hui Zhang were supported by the EPSRC grant EP/N018702/1 and Marco Palombo by the UKRI Future Leaders Fellowship MR/T020296/1.

Jan Morez is supported by a grant (ISLRA-2009) from the European Space Agency, by Belgian Science Policy Office-Prodex. Ben Jeurissen and Jan Sijbers received funding from the Research Foundation Flanders (FWO Vlaanderen: 12M3119N; G0D7216N).

Maryam Afzali and Derek K Jones are supported by a Wellcome Trust Investigator Award (096646/Z/11/Z) and DKJ by a Wellcome Trust Strategic Award (104943/Z/14/Z).

Tomasz Pieciak acknowledges the Polish National Agency for Academic Exchange for grant PN/BEK/2019/1/00421 under the Bekker programme and the Ministry of Science and Higher Education (Poland) under the scholarship for outstanding young scientists.

Fabian Bogusz acknowledges AGH Science and Technology, Poland (16.16.120.773).

Evren Özarslan is financially supported by Linköping University (LiU) Center for Industrial Information Technology (CENIIT), LiU Cancer, VINNOVA/ITEA3 17021 IMPACT, Analytic Imaging Diagnostic Arena (AIDA), and the Swedish Foundation for Strategic Research (RMX18-0056).

Andrada Ianus’ work received the support of a fellowship from “la Caixa” Foundation (ID 100010434) and from the European Union’s Horizon 2020 research and innovation programme under the Marie Skłodowska-Curie grant agreement No 847648, fellowship code LCF/BQ/PI20/11760029.

Santiago Aja-Fernandez acknowledges the “Ministerio de Ciencia e Innovación” of Spain for research grant RTI2018-094569-B-I00.

## Appendix

### Tensor-based models

- DTI: The diffusion tensor imaging method was fitted with a linear least squares procedure to determine the diffusion tensor (6 parameters) and the average non-weighted signal (1 parameter).
- DKI: The diffusion kurtosis imaging extends the DTI method to account for restricted diffusion. It was fitted with a weighted least squares procedure using ExploreDTI to determine 22 parameters: 6 for the diffusion tensor, 15 for the kurtosis tensor and the non-weighted signal. No additional constraints were considered in this fit.
- DKI+Offset: The DKI model was extended to accommodate an additional degree of freedom modelling a positive constant bias in the signal due to, for example, Rician noise. The 23 free parameters of this model were fitted with a non-linear least squares procedure implemented in MATLAB, constraining a monotonic signal decay and enforcing both the diffusion and kurtosis tensor to be positive definite.
- DTD-cov: The diffusion tensor distribution (DTD) method describes the diffusion signal as the sum of a distribution of microscopic tensors. The 28 parameters of the fourth order covariance tensor method were fitted to the data with a non-linear least squares procedure implemented in MATLAB, constraining a monotonic signal decay and enforcing both the diffusion and kurtosis tensor to be positive definite.
- DTD-cov+Offset: The DTD-cov method was extended with one additional degree of freedom modelling a positive constant bias in the signal due to, for example, Rician noise. The 29 free parameters of this model were fit with a non-linear least squares procedure implemented in MATLAB, constraining a monotonic signal decay and enforcing both the diffusion and kurtosis tensor to be positived definite.

### Multi-compartment models

- Ball&Stick: originally proposed from Behrens and colleagues, this model consists of two compartments: a stick (impermeable cylinder with zero radius) to model anisotropic restricted intra-cellular diffusion, and a ball to model isotropic hindered extra-cellular diffusion. The model was implemented in Python using the Dmipy package, and its 4 parameters fitted to the data using a two stages procedure consisting of an initial grid search, followed by a constrained non-linear fit procedure based on a limited-memory quasi-Newton method.
- Ball&Racket: this model is an extension of the Ball&Stick that explicitly takes into account fanning configurations. The 7 parameters of the model were fitted to the data using the same procedure described for the Ball&Stick model.
- NODDI-Watson: originally introduce from Zhang et al., this model accounts for intra-cellular diffusion modelled as a tensor convolved with a Watson distribution to account for axonal dispersion, an extracellular compartment modelled with a Zeppelin, and an isotropic free water component to account for partial volume with the cerebrospinal fluid. The volumes of the intracellular and extracellular compartments are linked with a tortuosity principle, and the parallel diffusivity of the tensor is set to 1.7×10^-3^mm^2^/s. The 5 parameters of the model were fitted to the data using the same procedure described for the Ball&Stick model.
- NODDI-Bingham: this model extends the NODDI-Watson model to account for asymmetric fiber dispersion using a Bingham distribution. The 7 parameters of the model were fitted to the data using the same procedure described for the Ball&Stick model.
- SMT: The spherical mean technique (SMT) model provides estimates of neurite density and of the intrinsic tissue diffusivity unconfounded by fibre crossings and orientation dispersion. The 51 parameters of the model were fitted to the data using the same procedure described for the Ball&Stick model.
- NODDI-SMT: This is a reformulation of the NODDI-Watson model using the SMT technique. The 50 parameters of the model were fitted to the data using the same procedure described for the Ball&Stick model.
- MCMDI: this model describes intra-cellular diffusion with a stick, and extra-cellular diffusion with a Zeppelin. The SMT technique is used to achieve invariance to fibre crossing and orientation dispersion. The 50 parameters of the model were fitted to the data using the same procedure described for the Ball&Stick model.
- ActiveAx: introduced from Dyrby and colleagues, this model describes intra-cellular diffusion as a cylinder with finite radius, extracellular diffusion as a zeppelin, and accounts for isotropic contamination due to cerebrospinal fluid. The 7 parameters of the model were fitted to the data using the same procedure described for the Ball&Stick model.

### Parametric representations

- SHORE: The method is based on the original simple harmonic oscillator reconstruction (SHORE) [1]. SHORE with optimized reconstruction was tested at different orders of 6, 8 and up to 12. However, the best results or lower errors were determined to be at either order 6 or 8. The 50 parameters of the model were fitted to the data using a linear least-squares approach.
- MAP-MRI: Mean Apparent Propagator Magnetic Resonance Imaging (MAP-MRI) is a linear representation of the diffusion signal that uses a 3D generalization of the SHORE basis. The 95 parameters of the method were fitted using a penalized least-squares procedure with generalized cross validation implemented in Dmipy.
- MAP-MRI+Reg: This submission used the Laplacian-regularized MAP-MRI method of order 8 implemented in the Dipy software library with no positivity constraint in the propagator and a regularization weight of 0.47 to fit the 95 free parameters of the method.

### Deep-learning methods

- NeuralNet: A fully connected neural network with a single hidden layer of 50 neurons and using sigmoid activation functions was trained to predict the unprovided signal amplitudes for each measurement independently. The 50 parameters of the network were optimized based on the mean squared error of the predictions using the ADAM algorithm with a learning rate of 0.005 over 20000 epochs. For SDE-MS and SDE-GRID the normalized components of the gradient (3 values) and the b-value were provided as inputs to the network. For the DDE and DODE acquisitions, the gradient strength, the normalized components of the two gradients (6 values), the b-value, and the components of the b-matrix (6 values) were concatenated into one input vector of length 14. • NeuralNet+Reinf: A fully connected neural network with reinforcement learning. The authors adopted a neural architecture search (NAS) to identify the optimal 7-layer perceptron model for dMRI signal prediction with either 8. 16, 32, 64 or 128 nodes per layer. The free parameters of the network ranged between 56 and 896, and their values was optimized with the ADAM method using an initial learning rate equal to 0.01 and 200 training epochs.

**Table A1:**
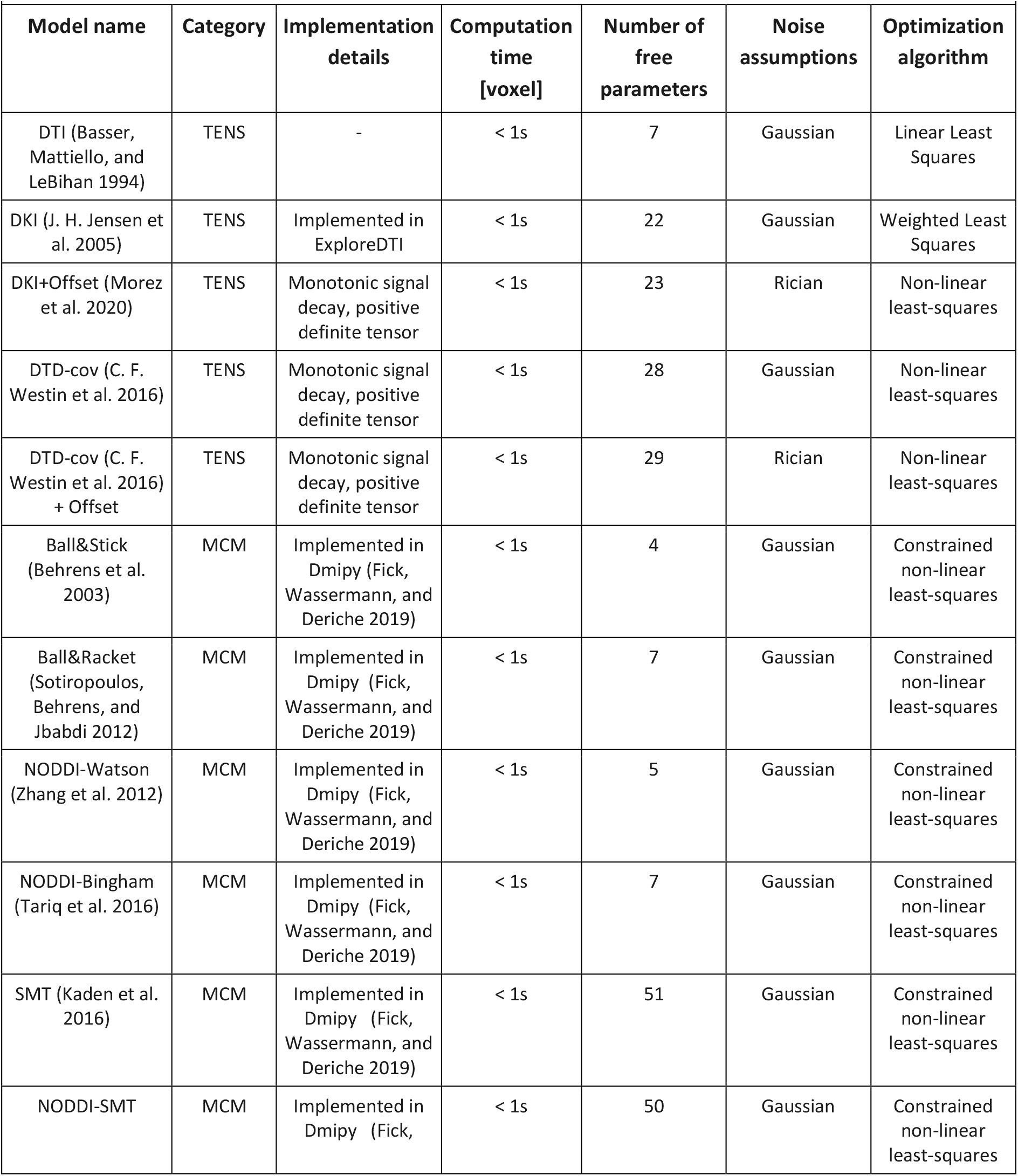
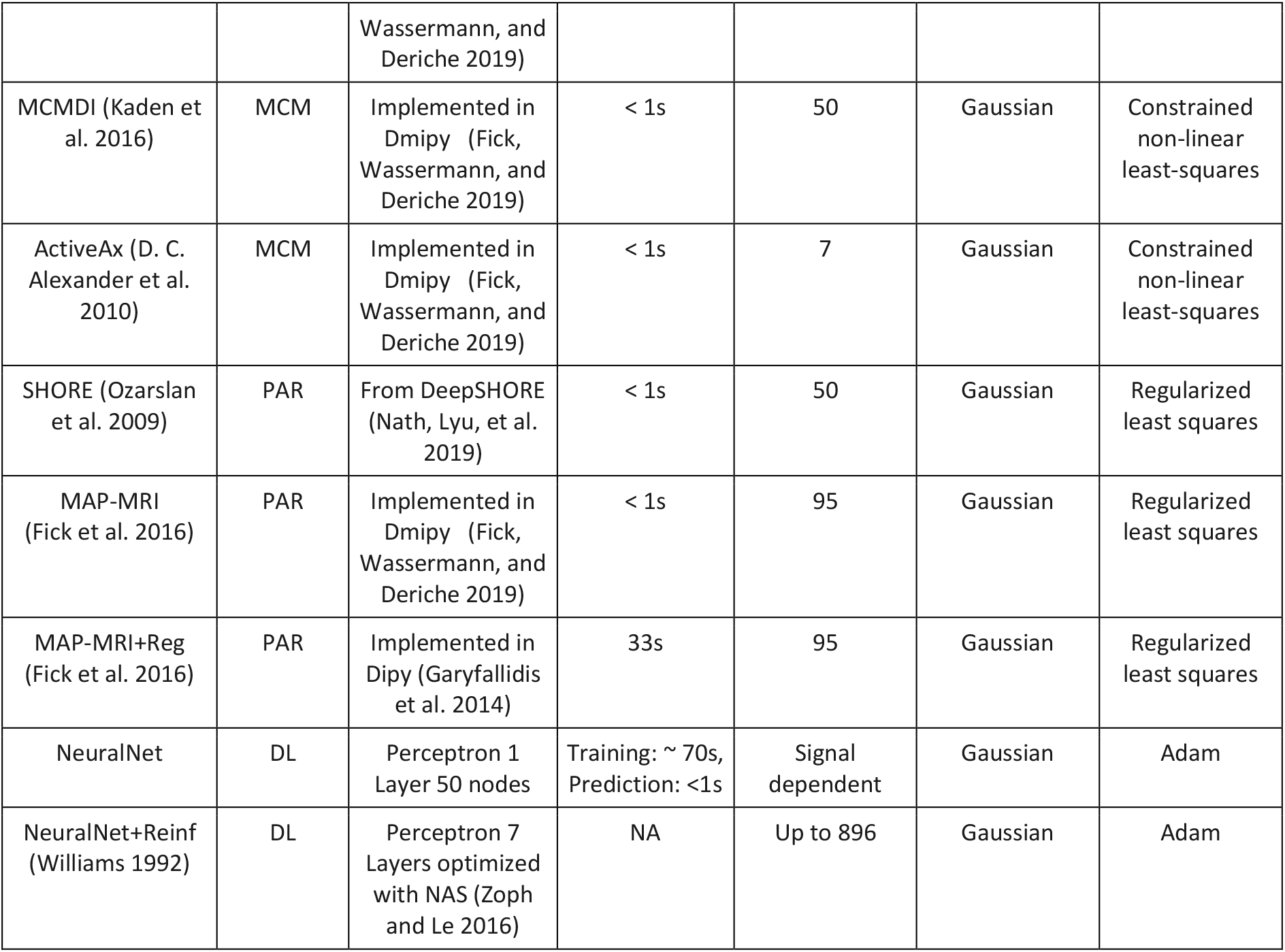
The valid signal predictions submitted to the MEMENTO challenge. For each method, we report the acronym and the main reference, the “category”, special notes on the fit procedure, and the data it has been applied to. The following predictions were subdivided in the following categories: tensor-based (TENS), multi-compartment model (MCM), parametric representation (PAR), deep learning-based (DL).

## Notes

### Competing Interest Statement

The authors have declared no competing interest.

### Summary of Updates

Updated after revision round

