## supplementary material for "On the generalizability of diffusion MRI signal representations across acquisition parameters, sequences and tissue types: chronicles of the MEMENTO challenge"

| **Table S1:** The mean squared error determined for each prediction of signals 1-5 with SDE-MS, SDE-GRID, DDE and DODE. The predictions with the lowest MSE are highlighted in BOLD. | | | | | | |
| --- | --- | --- | --- | --- | --- | --- |
| **Mean squared error (MSE)** | | | | | | |
| **Method** | **Signal 1** | **Signal 2** | **Signal 3** | **Signal 4** | **Signal 5** | **Average** |
| **SDE-MS** | | | | | | |
| ActiveAx | 0,00286 | 0,0041 | 0,00552 | 0,00842 | 0,00884 | 0,00595 ± 0,00235 |
| Ball&Racket | 0,00356 | 0,00221 | 0,0032 | **0,00243** | 0,00256 | 0,00279 ± 0,00051 |
| Ball&Stick | 0,00359 | 0,00563 | 0,00965 | 0,00295 | 0,00284 | 0,00493 ± 0,00256 |
| MAP-MRI+Reg | 0,0026 | 0,00186 | 0,00253 | 0,00245 | 0,00272 | 0,00243 ± 0,00030 |
| MCMDI | 0,0036 | 0,00277 | 0,00301 | 0,00566 | 0,00668 | 0,00434 ± 0,00155 |
| NeuralNet | 0,00336 | 0,00198 | 0,0025 | 0,00251 | **0,00164** | 0,00240 ± 0,00058 |
| NODDI-Watson | 0,00359 | 0,00339 | 0,00489 | 0,00346 | 0,00412 | 0,00389 ± 0,00056 |
| NODDI-Bingham | 0,00357 | 0,002 | 0,00313 | 0,00346 | 0,00408 | 0,00325 ± 0,00069 |
| SMT | 0,00442 | 0,00351 | 0,00484 | 0,00322 | 0,00279 | 0,00376 ± 0,00076 |
| NODDI-SMT | 0,0058 | 0,00476 | 0,00502 | 0,0062 | 0,00645 | 0,00565 ± 0,00066 |
| SHORE-worst | 0,01681 | 0,01522 | 0,01906 | 0,00578 | 0,00493 | 0,01236 ± 0,00585 |
| SHORE-best | **0,00249** | 0,00203 | 0,00285 | 0,00257 | 0,00275 | 0,00254 ± 0,00028 |
| MAP-MRI | 0,00291 | **0,00183** | **0,00238** | 0,00246 | 0,00222 | **0,00236 ± 0,00035** |
| DTI | 0,00804 | 0,0086 | 0,01128 | 0,00406 | 0,00454 | 0,00730 ± 0,00269 |
| NN+Reinf | 0,01779 | 0,00658 | 0,00639 | 0,00313 | 0,00278 | 0,00733 ± 0,00546 |
| DKI+Offset-worst | 0,00284 | 0,00198 | 0,0024 | 0,00245 | 0,00265 | 0,00246 ± 0,00029 |
| DKI+Offset-best | 0,00264 | 0,00195 | 0,00255 | 0,00245 | 0,0025 | 0,00242 ± 0,00024 |
| DKI | 0,00299 | 0,00211 | 0,00332 | 0,00245 | 0,00249 | 0,00267 ± 0,00043 |
| Minimum | 0,00249 | 0,00183 | 0,00238 | 0,00243 | 0,00164 | 0,00215 ± 0,00035 |
| **SDE-GRID** | | | | | | |
| MAP-MRI+Reg | 0,00321 | 0,00224 | **0,00238** | **0,00215** | **0,00301** | **0,00260 ± 0,00043** |
| NeuralNet-best | 0,0031 | 0,00231 | 0,00246 | 0,0023 | 0,00317 | 0,00267 ± 0,00039 |
| NeuralNet-worst | 0,00317 | 0,00218 | 0,00249 | 0,00241 | 0,00316 | 0,00268 ± 0,00041 |
| SHORE-best | 0,00329 | 0,00246 | 0,00256 | 0,00226 | 0,00311 | 0,00274 ± 0,00040 |
| SHORE-worst | 0,00324 | 0,0024 | 0,00246 | 0,00222 | 0,00312 | 0,00269 ± 0,00041 |
| NN+Reinf-worst | 0,01341 | 0,00715 | 0,00536 | 0,00383 | 0,00315 | 0,00658 ± 0,00368 |
| NN+Reinf-best | 0,01413 | 0,00632 | 0,00536 | 0,00383 | 0,00315 | 0,00656 ± 0,00395 |
| DKI+Offset-best | **0,00301** | **0,00206** | 0,00266 | 0,00216 | 0,00321 | 0,00262 ± 0,00045 |
| DKI+Offset-worst | 0,0048 | 0,00326 | 0,00328 | 0,00227 | 0,00319 | 0,00336 ± 0,00081 |
| Minimum | 0,00301 | 0,00206 | 0,00238 | 0,00215 | 0,00301 | 0,00252 ± 0,00041 |
| **DDE** | | | | | | |
| NeuralNet-worst | 0,00094 | 0,0015 | 0,00282 | 0,00429 | 0,00222 | 0,00235 ± 0,00116 |
| NeuralNet-best | 0,00057 | 0,00138 | 0,00301 | 0,0034 | 0,00103 | 0,00188 ± 0,00112 |
| SHORE-best | 0,00102 | 0,00215 | 0,00255 | 0,00125 | 0,00217 | 0,00183 ± 0,00059 |
| SHORE-worst | 0,00132 | 0,00234 | 0,00255 | 0,00136 | 0,00231 | 0,00198 ± 0,00053 |
| DKI | 0,00214 | 0,00538 | 0,01939 | 0,00492 | 0,01368 | 0,00910 ± 0,00643 |
| DTI | 0,00218 | 0,00245 | 0,00181 | 0,00149 | 0,00201 | 0,00199 ± 0,00033 |
| NN+Reinf-worst | 0,01892 | 0,01488 | 0,01693 | 0,0046 | 0,0085 | 0,01277 ± 0,00538 |
| NN+Reinf-best | 0,00078 | 0,00203 | 0,0011 | 0,001 | **0,00037** | 0,00106 ± 0,00055 |
| DTD-cov | 0,00059 | 0,00108 | **0,00068** | 0,00094 | 0,00041 | 0,00074 ± 0,00024 |
| DTD-cov+Offset | **0,00057** | **0,00105** | 0,00069 | **0,00089** | 0,0004 | **0,00072 ± 0,00023** |
| DKI+Offset-best | 0,00072 | 0,00148 | 0,00311 | 0,00118 | 0,00237 | 0,00177 ± 0,00086 |
| DKI+Offset-worst | 0,00082 | 0,00182 | 0,00352 | 0,0016 | 0,00286 | 0,00212 ± 0,00095 |
| Minimum | 0,00057 | 0,00105 | 0,00068 | 0,00089 | 0,00037 | 0,00071 ± 0,00024 |
| **DODE** | | | | | | |
| NeuralNet-best | **0,00044** | **0,00134** | **0,00069** | **0,00075** | **0,0003** | **0,00070 ± 0,00036** |
| NeuralNet-worst | 0,00114 | 0,0017 | 0,00094 | 0,00087 | 0,00035 | 0,00100 ± 0,00044 |
| SHORE-worst | 0,00402 | 0,0035 | 0,00249 | 0,00168 | 0,00372 | 0,00308 ± 0,00087 |
| SHORE-best | 0,00395 | 0,00355 | 0,00241 | 0,00173 | 0,00373 | 0,00307 ± 0,00086 |
| DKI | 0,00603 | 0,00757 | 0,01827 | 0,00591 | 0,0162 | 0,01080 ± 0,00533 |
| DTI | 0,0055 | 0,0045 | 0,00224 | 0,00217 | 0,00372 | 0,00363 ± 0,00129 |
| NN+Reinf-worst | 0,03331 | 0,02334 | 0,02467 | 0,03702 | 0,02919 | 0,02951 ± 0,00515 |
| NN+Reinf-best | 0,00287 | 0,0043 | 0,00142 | 0,00169 | 0,03058 | 0,00817 ± 0,01125 |
| DTD-cov | 0,00452 | 0,00339 | 0,00116 | 0,00148 | 0,0025 | 0,00261 ± 0,00124 |
| DTD-cov+Offset | 0,00435 | 0,00335 | 0,00117 | 0,00148 | 0,0025 | 0,00257 ± 0,00118 |
| Minimum | 0,00044 | 0,00134 | 0,00069 | 0,00075 | 0,0003 | 0,00070 ± 0,00036 |

| **Table S2:** The residual variance determined for each prediction of signals 1-5 with SDE-MS, SDE-GRID, DDE and DODE. The predictions with the lowest variance are highlighted in BOLD. | | | | | | |
| --- | --- | --- | --- | --- | --- | --- |
| **Residual variance** | | | | | | |
| **Method** | **Signal 1** | **Signal 2** | **Signal 3** | **Signal 4** | **Signal 5** | **Average** |
| **SDE-MS** | | | | | | |
| ActiveAx | 0,00283 | 0,00379 | 0,00502 | 0,00786 | 0,00704 | 0,00531 ± 0,00190 |
| Ball&Racket | 0,00356 | 0,00209 | 0,00301 | **0,00241** | 0,00207 | 0,00263 ± 0,00058 |
| Ball&Stick | 0,00357 | 0,00557 | 0,00952 | 0,00292 | 0,00267 | 0,00485 ± 0,00255 |
| MAP-MRI+Reg | 0,00259 | **0,00179** | 0,00243 | 0,00245 | 0,00207 | **0,00227 ± 0,00029** |
| MCMDI | 0,0036 | 0,00273 | 0,00289 | 0,00477 | 0,00495 | 0,00379 ± 0,00092 |
| NeuralNet | 0,00304 | 0,00197 | 0,00249 | 0,00251 | **0,0016** | 0,00232 ± 0,00049 |
| NODDI-Watson | 0,00357 | 0,00332 | 0,00478 | 0,00324 | 0,00294 | 0,00357 ± 0,00064 |
| NODDI-Bingham | 0,00355 | 0,00194 | 0,00302 | 0,00325 | 0,00292 | 0,00294 ± 0,00054 |
| SMT | 0,00409 | 0,00337 | 0,00482 | 0,0031 | 0,00278 | 0,00363 ± 0,00073 |
| NODDI-SMT | 0,00559 | 0,00452 | 0,00494 | 0,00618 | 0,00592 | 0,00543 ± 0,00062 |
| SHORE-worst | 0,01387 | 0,01129 | 0,01325 | 0,00464 | 0,00368 | 0,00935 ± 0,00433 |
| SHORE-best | **0,00249** | 0,00201 | 0,00265 | 0,00256 | 0,00266 | 0,00247 ± 0,00024 |
| MAP-MRI | 0,00288 | 0,00183 | **0,00234** | 0,00246 | 0,00214 | 0,00233 ± 0,00035 |
| DTI | 0,00677 | 0,00729 | 0,00916 | 0,00365 | 0,00404 | 0,00618 ± 0,00207 |
| NN+Reinf | 0,01734 | 0,00656 | 0,00575 | 0,00305 | 0,0021 | 0,00696 ± 0,00545 |
| DKI+Offset-worst | 0,00277 | 0,00196 | 0,00239 | 0,00245 | 0,00256 | 0,00243 ± 0,00027 |
| DKI+Offset-best | 0,00263 | 0,0019 | 0,00254 | 0,00245 | 0,00246 | 0,00240 ± 0,00026 |
| DKI | 0,00299 | 0,00211 | 0,00315 | 0,00245 | 0,00245 | 0,00263 ± 0,00038 |
| Minimum | 0,00249 | 0,00179 | 0,00234 | 0,00241 | 0,0016 | 0,00213 ± 0,00036 |
| **SDE-GRID** | | | | | | |
| MAP-MRI+Reg | 0,00321 | 0,00223 | **0,00238** | **0,00215** | **0,00301** | **0,00260 ± 0,00043** |
| NeuralNet-best | 0,0031 | 0,00229 | 0,00245 | 0,00227 | 0,00312 | 0,00265 ± 0,00038 |
| NeuralNet-worst | 0,00318 | 0,00218 | 0,00245 | 0,00229 | 0,0031 | 0,00264 ± 0,00042 |
| SHORE-best | 0,00323 | 0,00238 | 0,00255 | 0,00226 | 0,00311 | 0,00271 ± 0,00039 |
| SHORE-worst | 0,00319 | 0,00233 | 0,00246 | 0,00222 | 0,00312 | 0,00266 ± 0,00041 |
| NN+Reinf-worst | 0,01305 | 0,00591 | 0,00527 | 0,00303 | 0,00312 | 0,00608 ± 0,00367 |
| NN+Reinf-best | 0,01343 | 0,00609 | 0,00527 | 0,00303 | 0,00312 | 0,00619 ± 0,00381 |
| DKI+Offset-best | **0,00301** | **0,00206** | 0,00266 | 0,00215 | 0,00314 | **0,00260 ± 0,00044** |
| DKI+Offset-worst | 0,00432 | 0,00299 | 0,00321 | 0,00227 | 0,00319 | 0,00320 ± 0,00066 |
| Minimum | 0,00301 | 0,00206 | 0,00238 | 0,00215 | 0,00301 | 0,00252 ± 0,00041 |
| **DDE** | | | | | | |
| NeuralNet-worst | 0,00051 | 0,00106 | 0,00124 | 0,00196 | 0,00081 | 0,00112 ± 0,00049 |
| NeuralNet-best | 0,00036 | 0,00125 | 0,00158 | 0,00192 | 0,00048 | 0,00112 ± 0,00061 |
| SHORE-best | 0,00046 | 0,00138 | 0,00248 | 0,00122 | 0,00208 | 0,00152 ± 0,00070 |
| SHORE-worst | 0,00055 | 0,00143 | 0,00243 | 0,00129 | 0,00212 | 0,00156 ± 0,00066 |
| DKI | 0,00199 | 0,00532 | 0,01928 | 0,00493 | 0,0137 | 0,00904 ± 0,00643 |
| DTI | 0,00104 | 0,0014 | 0,00144 | 0,00119 | 0,00142 | 0,00130 ± 0,00016 |
| NN+Reinf-worst | 0,00541 | 0,00634 | 0,00473 | 0,00176 | 0,00304 | 0,00426 ± 0,00165 |
| NN+Reinf-best | 0,00076 | 0,00169 | 0,0009 | 0,00098 | **0,00036** | 0,00094 ± 0,00043 |
| DTD-cov | 0,00041 | 0,00093 | **0,00065** | 0,00084 | **0,00036** | 0,00064 ± 0,00023 |
| DTD-cov+Offset | **0,00034** | **0,00089** | 0,00068 | **0,00081** | 0,00039 | **0,00062 ± 0,00022** |
| DKI+Offset-best | 0,0005 | 0,00132 | 0,00311 | 0,00115 | 0,00233 | 0,00168 ± 0,00092 |
| DKI+Offset-worst | 0,0006 | 0,0017 | 0,00351 | 0,00152 | 0,00282 | 0,00203 ± 0,00102 |
| Minimum | 0,00034 | 0,00089 | 0,00065 | 0,00081 | 0,00036 | 0,00061 ± 0,00023 |
| **DODE** | | | | | | |
| NeuralNet-best | **0,00037** | **0,00134** | **0,00067** | **0,00075** | **0,00029** | **0,00068 ± 0,00037** |
| NeuralNet-worst | 0,0009 | 0,0017 | 0,00078 | 0,00082 | 0,00032 | 0,00090 ± 0,00045 |
| SHORE-worst | 0,00206 | 0,0024 | 0,00204 | 0,00144 | 0,00254 | 0,00210 ± 0,00038 |
| SHORE-best | 0,00246 | 0,0027 | 0,00216 | 0,00155 | 0,00291 | 0,00236 ± 0,00047 |
| DKI | 0,00301 | 0,0051 | 0,01697 | 0,00499 | 0,01398 | 0,00881 ± 0,00557 |
| DTI | 0,00461 | 0,004 | 0,00223 | 0,00207 | 0,00333 | 0,00325 ± 0,00099 |
| NN+Reinf-worst | 0,00259 | 0,00322 | 0,00199 | 0,00358 | 0,00224 | 0,00272 ± 0,00059 |
| NN+Reinf-best | 0,00225 | 0,00429 | 0,00137 | 0,00093 | 0,02051 | 0,00587 ± 0,00741 |
| DTD-cov | 0,00156 | 0,00149 | 0,00077 | 0,00106 | 0,00091 | 0,00116 ± 0,00031 |
| DTD-cov+Offset | 0,00158 | 0,00149 | 0,00072 | 0,00106 | 0,00083 | 0,00114 ± 0,00034 |
| Minimum | 0,00037 | 0,00134 | 0,00067 | 0,00075 | 0,00029 | 0,00068 ± 0,00037 |

| **Table S3:** The Bayesian information criteria determined for each prediction of signals 1-5 with SDE-MS, SDE-GRID, DDE and DODE. The predictions with the lowest BIC are highlighted in BOLD. | | | | | | |
| --- | --- | --- | --- | --- | --- | --- |
| **Bayesian information criteria (BIC)** | | | | | | |
| **Method** | **Signal 1** | **Signal 2** | **Signal 3** | **Signal 4** | **Signal 5** | **Average** |
| **SDE-MS** | | | | | | |
| ActiveAx | -14555,77 | -13661,36 | -12915,48 | -11864,29 | -11742,38 | -12947,86 ± 1069,84 |
| Ball&Racket | -14014,80 | -15198,07 | -14277,17 | **-14968,48** | -14835,79 | -14658,86 ± 442,36 |
| Ball&Stick | -14017,25 | -12893,05 | -11546,30 | -14506,31 | -14602,33 | -13513,05 ± 1155,84 |
| MAP-MRI+Reg | -14105,84 | -14938,97 | -14178,29 | -14252,11 | -13994,62 | -14293,97 ± 333,52 |
| MCMDI | -13645,78 | -14304,22 | -14092,34 | -12517,16 | -12105,32 | -13332,96 ± 870,69 |
| NeuralNet | 10087,65 | 8768,39 | 9342,20 | 9359,67 | 8288,47 | 9169,28 ± 607,63 |
| NODDI-Watson | -14003,54 | -14147,98 | -13237,79 | -14100,92 | -13661,24 | -13830,29 ± 341,71 |
| NODDI-Bingham | -14006,14 | **-15446,09** | -14334,71 | -14084,26 | -13674,80 | -14309,20 ± 606,31 |
| SMT | -13126,93 | -13705,70 | -12902,27 | -13917,33 | -14276,78 | -13585,80 ± 505,87 |
| NODDI-SMT | -12459,95 | -12952,68 | -12819,52 | -12291,39 | -12191,68 | -12543,05 ± 295,95 |
| SHORE-worst | -9802,77 | -10050,33 | -9489,07 | -12467,97 | -12863,01 | -10934,63 ± 1429,86 |
| SHORE-best | -14572,34 | -15080,75 | -14227,56 | -14486,55 | -14323,88 | -14538,21 ± 296,86 |
| MAP-MRI | -14174,89 | -15332,59 | -14667,57 | -14590,12 | **-14847,96** | -14722,62 ± 376,40 |
| DTI | -11979,98 | -11811,22 | -11134,53 | -13683,37 | -13407,66 | -12403,35 ± 978,45 |
| NN+Reinf | 221581,48 | 219101,77 | 219025,79 | 217247,76 | 216952,77 | 218781,92 ± 1655,22 |
| DKI+Offset-worst | -14392,91 | -15299,59 | **-14817,57** | -14766,30 | -14572,52 | -14769,78 ± 304,61 |
| DKI+Offset-best | **-14629,08** | -15388,16 | -14715,34 | -14816,45 | -14770,48 | **-14863,90 ± 269,46** |
| DKI | -14327,10 | -15195,76 | -14065,70 | -14824,27 | -14782,80 | -14639,13 ± 397,73 |
| Minimum | -14629,08 | -15446,09 | -14817,57 | -14968,48 | -14847,96 | -14941,84 ± 274,61 |
| **SDE-GRID** | | | | | | |
| MAP-MRI+Reg | -6045,20 | -6468,34 | -6395,95 | -6515,40 | -6120,25 | -6309,03 ± 190,14 |
| NeuralNet-best | 15185,63 | 14839,08 | 14913,23 | 14834,87 | 15211,77 | 14996,92 ± 167,30 |
| NeuralNet-worst | 24926,74 | 24488,83 | 24641,95 | 24602,34 | 24922,11 | 24716,39 ± 177,15 |
| SHORE-best | -6334,74 | -6675,34 | -6630,12 | -6773,81 | -6403,08 | -6563,42 ± 166,88 |
| SHORE-worst | -6352,64 | -6704,08 | **-6675,14** | -6797,22 | -6396,82 | -6585,18 ± 177,06 |
| NN+Reinf-worst | 125490,99 | 124755,16 | 124417,30 | 124023,75 | 123794,58 | 124496,36 ± 596,38 |
| NN+Reinf-best | 125552,05 | 124609,87 | 124417,30 | 124023,75 | 123794,58 | 124479,51 ± 607,96 |
| DKI+Offset-best | **-6476,60** | **-6915,95** | -6620,24 | -6863,69 | -6399,04 | **-6655,10 ± 205,03** |
| DKI+Offset-worst | -6085,50 | -6538,05 | -6530,33 | **-6959,26** | **-6563,97** | -6535,42 ± 276,74 |
| Minimum | -6476,60 | -6915,95 | -6675,14 | -6959,26 | -6563,97 | -6718,18 ± 190,38 |
| **DDE** | | | | | | |
| NeuralNet-worst | 21121,54 | 21344,72 | 21648,73 | 21850,49 | 21533,86 | 21499,87 ± 250,38 |
| NeuralNet-best | 7419,20 | 7845,51 | 8220,87 | 8279,00 | 7707,78 | 7894,47 ± 321,72 |
| SHORE-best | -2999,05 | -2639,45 | -2557,61 | -2899,96 | -2635,26 | -2746,26 ± 171,37 |
| SHORE-worst | -2873,62 | -2599,28 | -2557,06 | -2859,23 | -2604,14 | -2698,67 ± 138,03 |
| DKI | -2814,36 | -2372,48 | -1756,89 | -2414,99 | -1924,24 | -2256,59 ± 376,77 |
| DTI | -2898,20 | -2841,67 | -2986,92 | -3080,15 | -2937,99 | -2948,99 ± 81,07 |
| NN+Reinf-worst | 113946,77 | 113831,41 | 113893,32 | 113267,43 | 113562,71 | 113700,33 ± 253,58 |
| NN+Reinf-best | 112418,10 | 112875,54 | 112582,75 | 112533,14 | 112062,90 | 112494,48 ± 263,28 |
| DTD-cov | -3393,43 | -3107,20 | **-3330,15** | -3171,75 | -3571,09 | -3314,72 ± 164,72 |
| DTD-cov+Offset | **-3410,06** | **-3114,67** | -3316,98 | **-3195,29** | **-3572,03** | **-3321,81 ± 160,83** |
| DKI+Offset-best | -3286,25 | -2941,96 | -2586,34 | -3049,75 | -2716,56 | -2916,17 ± 246,62 |
| DKI+Offset-worst | -3268,33 | -2885,72 | -2569,55 | -2948,64 | -2669,27 | -2868,30 ± 243,16 |
| Minimum | -3410,06 | -3114,67 | -3330,15 | -3195,29 | -3572,03 | -3324,44 ± 160,84 |
| **DODE** | | | | | | |
| NeuralNet-best | 19488,64 | 20653,94 | 19960,37 | 20049,73 | 19109,09 | 19852,35 ± 524,96 |
| NeuralNet-worst | 5336,46 | 5756,60 | 5139,75 | 5054,55 | 4098,50 | 5077,17 ± 546,09 |
| SHORE-worst | -5390,25 | -5532,88 | -5889,79 | -6298,26 | -5469,61 | -5716,16 ± 337,52 |
| SHORE-best | -5409,02 | -5519,08 | -5921,44 | -6266,08 | -5468,98 | -5716,92 ± 328,28 |
| DKI | -5161,82 | -4925,55 | -4009,87 | -5182,82 | -4134,93 | -4683,00 ± 508,21 |
| DTI | -5362,18 | -5570,19 | -6295,23 | -6331,56 | -5770,18 | -5865,87 ± 387,69 |
| NN+Reinf-worst | 136214,35 | 135844,59 | 135902,21 | 136324,27 | 136077,11 | 136072,51 ± 181,37 |
| NN+Reinf-best | 133665,75 | 134084,14 | 132933,69 | 133114,50 | 136125,50 | 133984,72 ± 1145,12 |
| DTD-cov | -5420,16 | -5719,80 | **-6834,08** | **-6581,01** | **-6036,72** | -6118,35 ± 525,22 |
| DTD-cov+Offset | **-5454,03** | **-5724,79** | -6820,20 | -6572,20 | -6028,10 | **-6119,86 ± 510,47** |
| Minimum | -5454,03 | -5724,79 | -6834,08 | -6581,01 | -6036,72 | -6126,13 ± 515,54 |
